# Soil chemistry and soil history significantly structure oomycete communities in *Brassicaceae* crop rotations

**DOI:** 10.1101/2022.07.12.499733

**Authors:** Andrew J.C. Blakney, Luke D. Bainard, Marc St-Arnaud, Mohamed Hijri

## Abstract

Oomycetes are critically important soil microbial communities, especially for agriculture where they are responsible for major declines in yields. Unfortunately, oomycetes are vastly understudied compared to bacteria and fungi. As such, our understanding of how oomycete biodiversity and community structure varies through time in the soil remains poor. Soil history established by previous crops is one factor known to structure other soil microbes, but has not been investigated for its influence on oomycetes. In this study, we established three different soil histories in field trials; the following year these plots were planted with five different *Brassicaceae* crops. We hypothesized that the previously established soil histories would structure different oomycete communities, regardless of their current *Brassicaceae* crop host, in both the roots and rhizosphere. We used a nested-ITS amplicon strategy incorporated with MiSeq metabarcoding, where the sequencing data was used to infer amplicon sequence variants (ASVs) of the oomycetes present in each sample. This allowed us to determine the impact of different soil histories on the structure and biodiversity of the oomycete root and rhizosphere communities from the five different *Brassicaceae* crops. We found that each soil history structured distinct oomycete rhizosphere communities, regardless of different *Brassicaceae* crop hosts, while soil chemistry structured the oomycete communities more during a dry year. Interestingly, soil history appeared specific to oomycetes, but was less influential for bacterial communities previously identified from the same samples. These results advance our understanding of how different agricultural practices and inputs can alter edaphic factors to impact future oomycete communities. Examining how different soil histories endure and impact oomycete biodiversity will help clarify how these important communities may be assembled in agricultural soils.

**Highlights:**

- Crop rotations model how soil history impacts subsequent microbial communities
- *Brassicaceae* oilseed crops might mitigate pathogenic oomycetes
- Soil history significantly structures oomycete communities
- Oomycetes are significantly affected by soil chemistry
- *Brassicaceae* crop hosts weakly influence oomycete communities

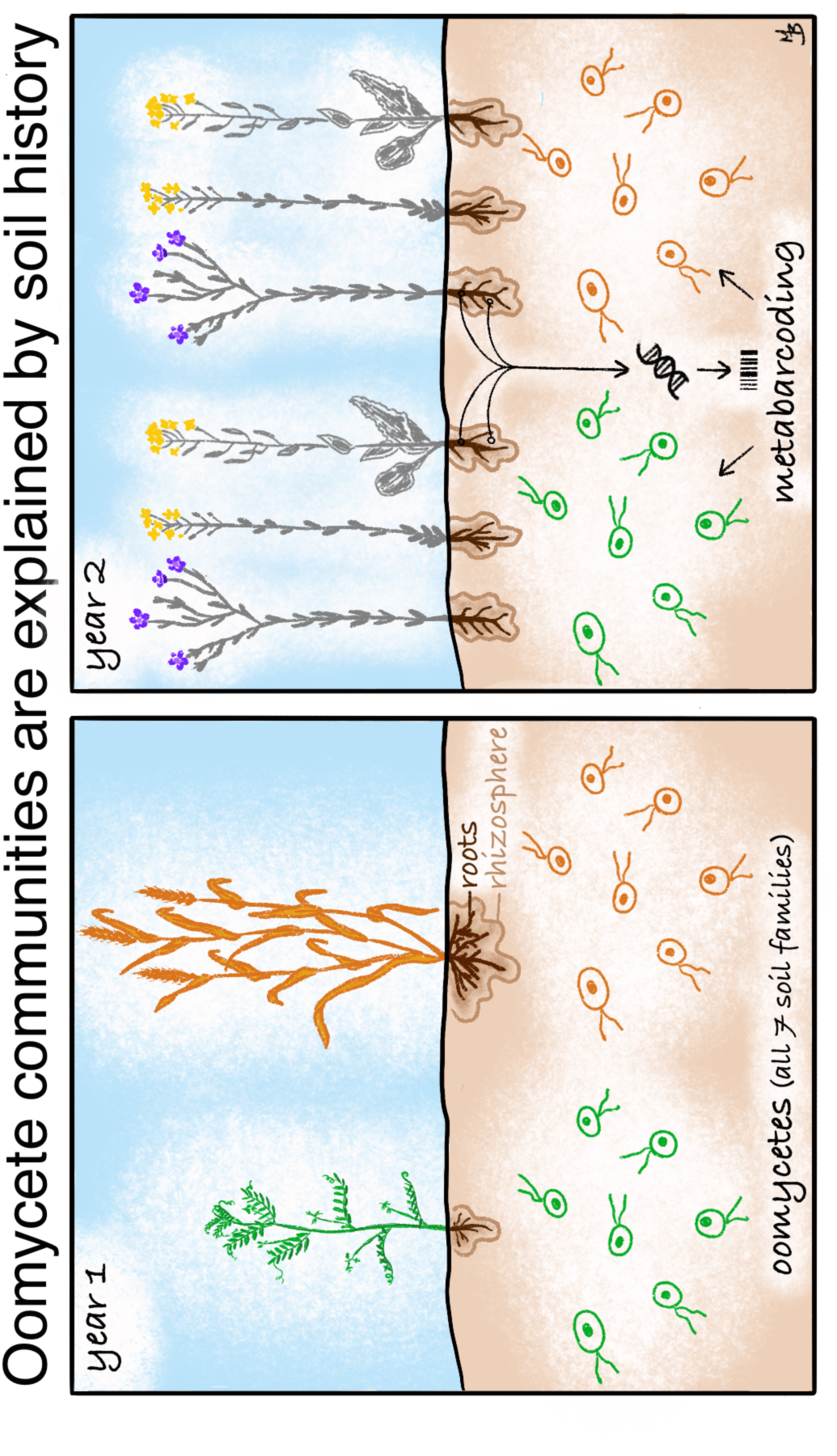

## 1. Introduction

Soil history established by previous plant-soil microbial communities’ condition future generations (Kaisermann *et al*., 2017; Bakker *et al*., 2021; Hannula *et al*., 2021; Blakney *et al*., 2022). These communities are subject to various biotic and abiotic factors—including the initial soil chemistry and microbes present, changes in the plant communities, and environmental extremes, such as droughts—and subsequently reflect them through a plant-soil microbial feedback process (Hwang *et al*., 2015; Yang *et al*., 2021; Liu *et al*., 2022). Plant hosts alter soil chemistry, first, through their capacity to uptake nutrients from the soil (Hu *et al*., 2021). Second, through rhizodeposition the host plant can vary the quantity and array of compounds released into the rhizosphere as required, thereby changing the soil chemistry (Lebeis *et al*., 2015; Korenblum *et al*., 2020; Kawasaki *et al*., 2021; Yu *et al*., 2021). Modifying their rhizodeposition profile also permits plants to actively tailor the structure of their microbial rhizosphere community in response to variable conditions and the plant’s needs (Lebeis *et al*., 2015; Korenblum *et al*., 2020; Kawasaki *et al*., 2021; Yu *et al*., 2021). For example, soil microbes increase the host plant’s access to nutrients (Richardson *et al*., 2009; Weidner *et al*., 2015; Yu *et al*., 2021), temper environmental change (Lau & Lennon, 2012), or stress (Marasco *et al*., 2012; Hou *et al*., 2021), and protect against pathogens (Mendes *et al*., 2011; Sikes *et al*., 2009). Soil bacterial communities in particular help integrate these diverse signals and modulate the plant’s responses (Castrillo *et al*., 2017; Hou *et al*., 2021).

As such, the plant-soil microbial community generates a reciprocal feedback process that incorporates various biotic and abiotic factors (Hwang *et al*., 2015; Yang *et al*., 2021; Liu *et al*., 2022), and impacts future plant-soil microbial generations and their composition (Kaisermann *et al*., 2017; Berendsen *et al*., 2018; Fitzpatrick *et al*., 2018). Thus, information from one plant-soil microbial community is transmitted through time to impact subsequent plant-microbial generations, i.e., that the soil history, also referred to as soil legacy, of previous plant-soil microbial communities condition future ones (Kaisermann *et al*., 2017; Bakker *et al*., 2021; Hannula *et al*., 2021). However, how different biotic and abiotic factors interact to establish soil histories are not well understood.

Drought, for example, is an increasingly important abiotic factor for establishing plant-soil microbial communities, as well as its growing impact on global agricultural production (Preece *et al*., 2019). On the Canadian Prairies, drought is a common experience. During the last event of the 2021 season, major crop production plunged nationally to a 15-year low due to severe drought conditions: wheat decreased by 38.5% to 21.7 M tonnes, while canola decreased by 35.4% to 12.6 M tonnes (Statistics Canada, November 2021). Water availability is a key determinant of plant nutrition and performance, such that their growth becomes restricted due to drought, as they are constrained in nutrient uptake from the soil (Fitzpatrick *et al*., 2018). Soil moisture is also critical for microbial communities (reviewed by Schimel *et al*., 2007); bacteria depend on water for nutrient diffusion and mobility (Preece *et al*., 2019). Soil moisture is also a key promotor of phytopathogenic oomycete growth and dispersion (Hwang *et al*., 2015; Rojas *et al*., 2017; Karppinen *et al*., 2020). As such, there is interest in investigating how soil history and plant-microbial communities interact under drought conditions in an agricultural setting.

Crop rotations, complete with their agricultural inputs, model how a previous crop plant establishes a soil history by altering the biotic and abiotic soil conditions for future plant-microbial communities (Hwang *et al*., 2015; Yang *et al*., 2021; Liu *et al*., 2022). For example, when lentils, or other legumes, are introduced into a rotation they shift the soil microbial community, which establishes more bioavailable nitrogen and moisture in the soil (O’Donovon *et al*., 2014; Bazghaleh *et al*., 2016; Hamel *et al*., 2018; Yang *et al*., 2021). These benefits translate to the subsequent crops, which tend to have higher yields (O’Donovon *et al*., 2014; Hamel *et al*., 2018).

Beneficial soil histories have also been established by canola (cultivars of *Brassica napus*, *B*. *rapa*, or *B*. *juncea*) rotations, as they are thought to reduce the growth of cereal-specific pathogens. As such, cereals tend to have higher yields when they are planted after canola (Etesami & Alikhani, 2016; Yang *et al*., 2021). As demand for vegetable oil and biofuels increases, *Brassicaceae* oilseed-based rotations are increasingly common throughout the world, such as the frequent canola-wheat rotation in Canada (Yang *et al*., 2021). Increasing *Brassicaceae* oilseed crop diversity has been on-going in order to improve production by identifying and breeding crop plants better adapted to the drought stress of the Canadian Prairies, as well as cultivars that are more resistant to pathogens (Bailey-Serres et al., 2019; Hossain et al., 2019; Liu et al., 2019). Though extensive work has investigated how *Brassicaceae* oilseed rotations benefit the crop plants involved (recently reviewed by Yang *et al*., 2021), less is known how these crops and their agricultural inputs impact beneficial, or pathogenic soil microbial communities.

*Brassicaceae* oilseed crops are known hosts to many microbial pathogens, including fungi, such as *Fusarium* sp. *Leptosphaeria* sp. (blackleg), *Rhizoctonia solani*, *Sclerotinia sclerotiorum* (stem rot), and oomycetes, like *Pythium* sp., *Phytophthora* sp., *Albugo* sp. (staghead), and *Peronospora* and *Hyaloperonospora* spp. (downy mildews) (Canola Encyclopedia: Diseases, 2017; Maciá-Vicente *et al*., 2020). Although specific *Brassicaceae* plants have been fairly well studied as model patho-systems for particular oomycetes, they remain a relatively under-studied group of phytopathogens, despite an historically outsized impact on global agriculture (Cevik *et al*., 2019; Derevnina *et al*., 2016; Kamoun *et al*., 2015; Mohammed *et al*., 2019; Price *et al*., 2017).

A few high-throughput sequencing studies have yielded some clues concerning soil oomycete communities: Sapp *et al*. (2018) and Maciá-Vicente *et al*. (2020) both illustrated oomycete communities present in agricultural and natural *Brassicaceae* soil samples, respectively. They identified root and rhizosphere communities dominated by *Pythiales* (*Pythium* sp., or *Globisporangium* sp.), with minorities of *Peronosporales* (*Phytophthora* sp., *Peronospora* sp., and *Hyaloperonospora* sp.), and *Saprolegniales* (*Aphanomyces* sp.), though they indicated substantial difficulties in assigning taxa (Sapp *et al*., 2018; Maciá-Vicente *et al*., 2020). Taheri *et al*. (2017) yielded further insights by also identifying oomycete root and rhizosphere communities dominated by *Pythiales* from various agricultural pea fields across the Canadian Prairies. Beyond these examples, there is a lack of baseline knowledge concerning how oomycete communities are structured around diverse *Brassicaceae* oilseed crops.

Crop rotations and the soil histories they establish, highlight important factors to investigate in order to better understand oomycete community dynamics. First, different crops, along with their agricultural treatments, alter edaphic factors, which may have important consequences for oomycete communities. For example, legumes tend to retain soil moisture, which is a key factor for oomycete growth as discussed (Hwang *et al*., 2015; Rojas *et al*., 2017; Karppinen *et al*., 2020). Second, crop rotations are known to shift bacterial communities, which may interact to suppress or promote, oomycete communities (Löbmann *et al*., 2016). Both of these examples stress what impact crop rotations might have on the subsequent crop plant-oomycete community?

We took advantage of an existing agriculture experiment to investigate the impact of soil history established by the previous year’s crop, and the current *Brassicaceae* oilseed host crops, on the soil oomycete communities. Three crop histories were established by growing wheat, lentils, or being left fallow (Fig. S1A). The following year, each crop history plot was divided into five subplots and planted with a different *Brassicaceae* oilseed crop. At full flower, the root systems of these *Brassicaceae* host crops were harvested, and divided into root and rhizosphere compartments, from which environmental DNA was extracted. We hypothesized that the three soil histories established by the previous crops would structure different oomycete communities, regardless of their current *Brassicaceae* host, in both the roots and rhizosphere. We used a MiSeq metabarcoding approach to specifically target the oomycete communities for each root and rhizosphere sample, where the sequencing data was used to infer amplicon sequence variants (ASVs) to identify the oomycete biodiversity, and how this understudied group varies in agricultural soils.

## 2. Materials & Methods

### 2.1 Site and experimental design

A field experiment was conducted at the experimental farm of Agriculture and Agri-Food Canada’s Research and Development Centre, in Swift Current, Saskatchewan (50°15′N, 107°43′W). The site is located in the semi-arid region of the Canadian Prairies; according to the weather station of the research farm, the 2016 and 2017 growing seasons (May, June and July) had 328.4 mm and 55.0 mm of precipitation, respectively; compared to the 30-year average [1981-2010] of 169.2 mm. The daily temperature averages for the 2016 and 2017 seasons were 15.6°C and 15.9°C, respectively, while the 30-year average was 14.93°C. The farm is on a Brown Chernozem with a silty loam texture (46% sand, 32% silt, and 22% clay), and has been well-described previously by Liu *et al*., (2019 & 2020).

The experiment was established in a field previously growing spring wheat (*Triticum aestivum* cultivar AC Lillian). A two-phase cropping sequence—consisting of a Conditioning Phase the first year, and a Test Phase in the second year (Fig. S1)—was repeated in two field trials, Trial 1, 2015-2016, and Trial 2, 2016-2017, on adjacent sites (Fig. S1B & C). On each site, the experimental design was a split-plot replicated in four complete blocks. In the Conditioning Phase, three soil history treatments were randomly assigned to the main plots, consisting of spring wheat (*Triticum aestivum*, cv. AC Lillian), red lentil (*Lens culinaris* cv. CDC Maxim CL), or left fallow (Fig. S1). Thus, the Conditioning Phase established a soil history composed of either wheat, lentil, or fallow, plus their respective management plans as described below (Hossain *et al*., 2019; Liu *et al*., 2019; Blakney *et al*., 2022).

In the Test Phase, the 12 Conditioning Phase plots were each subdivided and five *Brassicaceae* oilseed crop species were randomly assigned to one of these five subplots (Fig. S1). The *Brassicaceae* crops seeded were Ethiopian mustard (*Brassica carinata* L., cv. ACC110), canola (*B*. *napus* L., cv. L252LL), oriental mustard (*B*. *juncea* L., cv. Cutlass), yellow mustard (*Sinapis alba* L., cv. Andante), and camelia (*Camelina sativa* L., cv. Midas). Therefore, the Test Phase established the *Brassicaceae* host plant-soil microbial community feedback, composed of the individual *Brassicaceae* genotypes, their soil microbial community, and their respective management plans, as described below (Hossain *et al*., 2019; Liu *et al*., 2019; Blakney *et al*., 2022). In total, each field trial had 60 subplots to sample (Fig. S1 & S2). For further details of this well-described experiment, its design, and treatments, see Hossain *et al*. (2019), Liu *et al*. (2019), and Blakney *et al*. (2022).

### 2.2 Crop rotation management and sampling

Crops in both field trials were grown and maintained according to standard management practices, as previously described by Hossain *et al*. (2019), Liu *et al*. (2019), and Blakney *et al*. (2022). A pre-seed ‘burn off’ herbicide treatment using glyphosate (Roundup, 900 g acid equivalent per hectare, a. e. ha^−1^) was applied to all plots each year to ensure a clean starting field prior to seeding. Lentil seeds were treated with a commercial rhizobium-based inoculant (TagTeam at 3.7 kg ha^−1^). Lentil and wheat were direct-seeded into wheat stubble from late April to mid-May, depending on the crop and year. The herbicides, glyphosate (Roundup, 900 g a. e. ha^−1^), Assure II (36 g active ingredient per hectare, a. i. ha^−1^), and Buctril M (560 g a.i. ha^−1^) were applied to the fallow, lentil, and wheat plots, respectively, for in-season weed control, while fungicides were only applied as needed. Soil tests were used to determine the rates of in-season nitrogen, phosphorus, and potassium application; no synthetic nitrogen fertilizer was applied to the lentil plots during the Conditioning Phase. Both lentil and wheat were harvested between late August and early October, depending on the crop and year.

The subsequent Test Phase *Brassicaceae* plant hosts were subjected to the same standard management practices as the Conditioning Phase, including pre-seed ‘burn off’, in-season herbicide and fungicide treatments as needed, and fertilized as recommended by soil tests (Table S1; Hossain *et al*. 2019, Liu *et al*. 2019, and Wang *et al*. 2020). Additionally, all *Brassicaceae* crops, except *B*. *napus*, were treated with Assure II mixed with Sure-Mix or Merge surfactant (0.5% v/v) for post-emergence grass control: Liberty (glufosinate, 593 g a.i. ha−1) was used for *B*. *napus*.

**Table 1.** The oomycete rhizosphere communities had more unique ASVs than the root communities of five *Brassicaceae* host plants in the Test Phase of a two-year crop rotation, harvested from two field trials (Trial 1, 2016; Trial 2, 2017) from Swift Current, SK. 17 656 076 raw reads were produced via Illumina’s MiSeq at Génome Québec, and processed through DADA2, where 8 068 013 reads were retained (ITS reads reported here) for ASV inference. A total of 1037 ASVs were identified across the entire dataset, which were filtered to 412 oomycete ASVs.

|  |  | Retained ITS Reads | ASV Occurance | ASV Abundance |
| --- | --- | --- | --- | --- |
| Trial 1 | Rhizosphere | 2 178 736<br>(36 312 ± 12 084 / sample) | 227<br>(69 ± 34 / sample) | 1 353 787<br>(22 563 ± 15 585 / sample) |
|  | Roots | 1 785 905<br>(29 765 ± 11 416 / sample) | 89<br>(63 ± 36 / sample) | 81 396<br>(1357 ± 4627 / sample) |
| Trial 2 | Rhizosphere | 2 355 829<br>(39 263 ± 14 105 / sample) | 218<br>(74 ± 40 / sample) | 1 237 771<br>(20 629 ± 16 799 / sample) |
|  | Roots | 1 747 543<br>(29 126 ± 11 252 / sample) | 43<br>(56 ± 39 / sample) | 1820<br>(33 ± 147 / sample) |

Test phase *Brassicaceae* plants were sampled in mid-late July at full flowering, i.e. when 50% of the flowers on the main raceme were in bloom, as described by the Canola Council of Canada (Canola Encyclopedia: Canola Growth Stages, 2017), where flowering corresponds to higher activity in rhizosphere microbial communities (Chaparro *et al*., 2014). Four plants from two different locations within each subplot were excavated and pooled together as a composite sample (Hossain *et al*., 2019; Liu *et al*., 2019; Wang *et al*., 2020; Blakney *et al*., 2022). In the field, each plant had its rhizosphere soil divided from the root material by gently scraping it off using bleach sterilized utensils into fresh collection trays. The roots were then gently washed three times with sterilized distilled water to remove any soil. Both the rhizosphere and root portions were immediately flash-frozen and stored in liquid nitrogen vapour shipping containers until stored in the lab at −80°C (Delavaux *et al*., 2020). Based on the sampling strategy, in this study we define the rhizosphere microbiome as the microbial community in the soil in close contact with the roots (Hannula *et al*., 2021), and the root microbiome as the microbial community attached to, and within, the roots (Berendsen *et al*., 2018). Two additional soil cores were sampled from each plot, pooled, and kept on ice in coolers. These samples were homogenized in the lab and sieved to remove rocks and roots. They were then used for soil chemistry analyses, including total carbon, nitrogen, pH, and micronutrients (see Wang *et al*., 2020 for details). Aerial portions of each harvested plant sample were retained to determine dry weight (Fig. S3).

### 2.3 DNA extraction from Test Phase *Brassicaceae* root and rhizosphere samples

Nucleic acids were extracted from Trial 1 Test Phase *Brassicaceae* samples, for both rhizosphere and root portions. First, all the root samples were ground in liquid nitrogen via sterile mortar and pestles (Fig. S2). Total DNA and RNA were extracted from ~1.5 g of rhizosphere soil using the RNA PowerSoil Kit with the DNA elution kit (Qiagen, Germany). DNA and RNA were extracted using ~0.03 g of roots using the DNeasy Plant DNA Extraction Kit, and RNeasy Plant Mini Kit (Qiagen, Canada), respectively, following the manufacturer’s instructions (see Wang *et al*. (2020) for use of the RNA samples). All remaining harvested material from Trial 1 and 2, as well as the extracted DNA from Trial 1, were kept at −80°C before being shipped to Université de Montréal’s Biodiversity Centre, Montréal (QC, Canada) on dry ice for further processing (Delavaux *et al*., 2020; Lay *et al*., 2018).

Total DNA was extracted from the Trial 2 Test Phase samples; ~500 mg of rhizosphere soil was used for the NucleoSpin Soil gDNA Extraction Kit (Macherey-Nagel, Germany), and ~130 mg of roots for the DNeasy Plant DNA Extraction Kit (Qiagen, Germany) (Lay *et al*., 2018). A no-template extraction negative control was used with both the root and rhizosphere extractions and included with the Test Phase samples (Fig. S2), to assess the influence of the extraction kits on our sequencing results, and the efficacy of our lab preparation. All 242 extracted DNA samples (60 plots x 2 parallel field trials x 2 compartments, rhizosphere and root, +2 no-template extraction control samples) were quantified using the Qubit dsDNA High Sensitivity Kit (Invitrogen, USA), and qualitatively evaluated by mixing ~2 μL of each sample with 1 μL of GelRed (Biotium), and running it on a 0.7 % agarose gel for 30 minutes at 150 V. The no-template extraction negative controls were confirmed to not contain DNA after extraction, where the detection limit was > 0.1 ng (Qubit, Invitrogen, USA). Samples were kept at −80°C (Bell *et al*., 2016; Delavaux *et al*., 2020).

### 2.4 Assembly of oomycete mock community

To assess potential bias caused by lab manipulations, sequencing and downstream bioinformatic processing, we assembled an oomycete mock community of known composition from twenty species with staggered copy numbers of the ITS1 region. To do so, we followed the method of Bakker (2018), beginning with generating a standard curve of the copy numbers of the ITS region from DNA extracted from *Pythium ultimum* strain 6358.Ba.B, generously provided by Dr. S. Chatterton (Agriculture and Agri-Food Canada, Lethbridge Research and Development Centre). We then used this standard curve to estimate the ITS copy number from other oomycete DNA samples (Table S3).

**Table 3.** PERMANOVA identified that the compartment and soil history were significant experimental factors in structuring the oomycete communities from the Test Phase of a two-year crop rotation, harvested in 2016 (Trial 1) and 2017 (Trial 2) from Swift Current, SK. *Brassicaceae* host crops were only significant in the Test Phase oomycete communities of Trial 1, while the *Brassicaceae* host ~ soil history interaction was never significant. The PERMANOVA was calculated using a Bray-Curtis distance matrix, with 9999 permutations.

|  | Trial 1 |  |  | Trial 2 |  |  |
| --- | --- | --- | --- | --- | --- | --- |
|  | F Model | R<sup>2</sup> | Pr (> F) | F Model | R<sup>2</sup> | Pr (> F) |
| Compartment<sup>a</sup> | 17.8018 | 0.12264 | **0.0001** | 8.3717 | 0.07508 | **0.0001** |
| Soil History | 2.1235 | 0.02926 | **0.0015** | 1.6600 | 0.02977 | **0.0018** |
| *Brassicaceae* Host | 1.8876 | 0.05202 | **0.0003** | 0.9364 | 0.03359 | 0.6464 |
| Compartment ~<br>Soil History | 1.3196 | 0.01818 | 0.0941 | 1.6202 | 0.02906 | **0.0020** |
| Compartment ~<br>*Brassicaceae* Host | 1.7741 | 0.04889 | **0.0008** | 1.0426 | 0.03740 | 0.3031 |
| Soil History ~<br>*Brassicaceae* Host | 1.0734 | 0.05916 | 0.2367 | 0.8299 | 0.05954 | 0.9822 |
| Compartment ~<br>Soil History ~<br>*Brassicaceae* Host | 0.9040 | 0.04982 | 0.7576 | 0.8778 | 0.06298 | 0.9235 |

a, Rhizosphere or roots
b, Fallow, lentil, or wheat
c, *Brassica carinata*, *B. napus*, *B. juncea*, *Sinapis alba*, or *Camelina sativa*

A ~1 kb fragment containing the entirety of the ITS 1 region, the 5.8S gene, and the ITS 2 region, was PCR amplified from *P*. *ultimum* 6358.Ba.B using the ITS4 and ITS6 primers (Sapkota & Nicolaisen, 2015; Table S2; Alpha DNA, Montréal, Canada). The ITS4/6 PCR reaction consisted of 11.5 μL dH_2_O, 5.0 μL of 10X buffer (Qiagen, Canada), 2.5 μL of 10 μM ITS4 and ITS6 primers (Alpha DNA, Montréal, Canada), 1.0 μL of dNTPs (Qiagen, Canada), 0.5 μL of *T*. *aq* polymerase (Qiagen, Canada), and 2 μL of a 1:10 dilution of the template, for a total volume of 25 μL. The PCR was run in an Eppendorf Mastercycle ProS (Mississauga, ON, Canada) thermocycler and consisted of an initial denaturation of 2 minutes at 95°C, followed by 30 cycles of 30 seconds denaturation at 95°C, 30 seconds annealing at 55°C, and 1 minute elongation at 72°C, before a final elongation of 5 minutes at 72°C (Sapkota & Nicolaisen, 2015). Four μL of the ITS PCR product was mixed with 1 μL of loading dye containing Gel Red (Biotium), and visualized on a 1% agarose gel after 60 minutes at 100 V. The amplified ITS fragment was quantified using the QuBit dsDNA High Sensitivity Kit (Invitrogen, USA), then serially diluted to 10^-9^, where 1 μL of each dilution was then used as template in a 10 μL qPCR reaction.

**Table 2.**
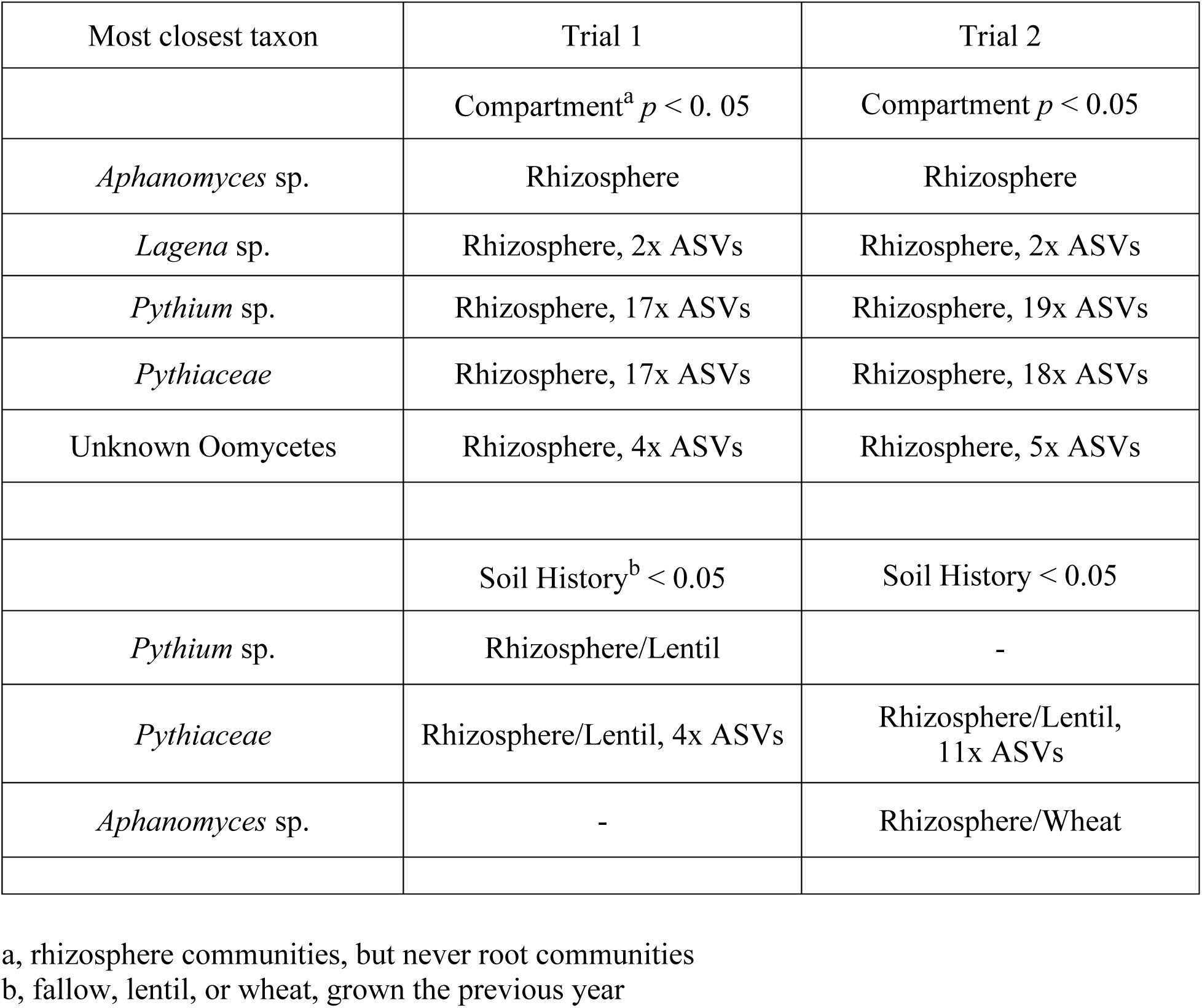
Indicator species were identified exclusively in the oomycete rhizosphere communities in the Test Phase of a two-year crop rotation, harvested in 2016 (Trial 1) and 2017 (Trial 2) from Swift Current, SK. In Trial 1, 41 ASVs were identified as indicator species in the Test Phase rhizosphere communities, while 45 ASVs were identified in Trial 2, notably from the same taxonomic groups. ASVs from the rhizosphere communities were also significant in specific soil histories established in the previous Conditioning Phase: in Trial 1, five ASVs were associated with the lentil soil history, while in Trial 2, 11 ASVs were again associated with the lentil soil history, and one ASV was associated with wheat soil history. No indicator species were identified for any of the five *Brassicaceae* host crops. Indicator species analysis relies on abundance and site specificity to statistically test each ASV, which we report here as *p* < 0.05, with a FDR correction.

| Most closest taxon | Trial 1 | Trial 2 |
| --- | --- | --- |
| | Compartment<sup>a</sup> $p < 0.05$ | Compartment $p < 0.05$ |
| *Aphanomyces* sp. | Rhizosphere | Rhizosphere |
| *Lagenia* sp. | Rhizosphere, 2x ASVs | Rhizosphere, 2x ASVs |
| *Pythium* sp. | Rhizosphere, 17x ASVs | Rhizosphere, 19x ASVs |
| *Pythiaceae* | Rhizosphere, 17x ASVs | Rhizosphere, 18x ASVs |
| Unknown Oomycetes | Rhizosphere, 4x ASVs | Rhizosphere, 5x ASVs |
| | Soil History<sup>b</sup> $p < 0.05$ | Soil History $p < 0.05$ |
| *Pythium* sp. | Rhizosphere/Lentil | - |
| *Pythiaceae* | Rhizosphere/Lentil, 4x ASVs | Rhizosphere/Lentil, 11x ASVs |
| *Aphanomyces* sp. | - | Rhizosphere/Wheat |

a, rhizosphere communities, but never root communities
b, fallow, lentil, or wheat, grown the previous year

The ITS1 region qPCR reactions amplified ~350 bp region using the oomycete-specific ITS6 forward and ITS7-a.e. reverse primers (Taheri *et al*., 2017). All qPCR reactions were set-up in triplicate in 96-well plates using the Freedom EVO100 robot (Tecan, Switzerland), with a triplicate no-template negative control included on each plate. The reactions consisted of 5.0 μL of Maxima SYBR Green/ROX qPCR Mix (ThermoFisher Scientific, Canada), 3.4 μL dH_2_O, 0.3 μL of 10 μM ITS6 and ITS7-a.e. primers (Alpha DNA, Montréal, Canada), and were run in a ViiA 7 Real-Time PCR System (ThermoFisher Scientific, Canada). The cycling conditions consisted of an initial denaturation of 2 minutes at 94°C, followed by 30 cycles of 30 seconds denaturation at 94°C, 30 seconds annealing at 59°C, and 1 minute elongation at 72°C, before a final elongation of 10 minutes at 72°C (Taheri *et al*., 2017). The number of ITS1 region copies present in the serially diluted standards were calculated using the formula (Godornes *et al*., 2007):

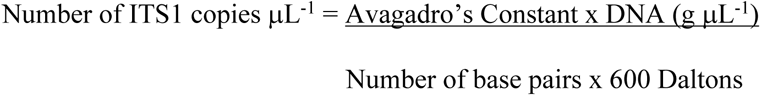

The standard curve was plotted, with an *R^2^* value of 0.9938 and an amplification efficiency of −3.389 (Fig. S4) falling within acceptable values (Fierer *et al*., 2005).

Copy numbers of the ITS1 region were then estimated for each oomycete sample (Table S3). One μL of a 1:10 dilution of extracted oomycete DNA was used as template for the ITS1 qPCR reaction, following the cycling conditions as described for generating the standard curve. Melt curves generated by 0.5°C increments at the end of the qPCR programme confirmed amplicon specificity. The mean cycle threshold was then calculated from the qPCR reactions for each oomycete sample, and the corresponding ITS1 copy number was estimated off the standard curve (Fig. S4; Bakker, 2018). Finally, the oomycete mock community was assembled with staggered ITS1 copy numbers, where different oomycete community members had different ranges of ITS1 copy numbers (Bakker, 2018).

### 2.5 ITS amplicon generation and sequencing to estimate the composition of the oomycete community

To estimate the composition of the oomycete communities in the rhizosphere and roots from the Test Phase *Brassicaceae* species, extracted DNA from all samples were used to prepare ITS amplicon libraries following Illumina’s MiSeq protocols. First, all DNA samples were diluted 1:10 into 96-well plates using the Freedom EVO100 robot (Tecan, Switzerland). To assess potential bias caused by lab manipulations, sequencing and downstream bioinformatic processing, we included a no-template DNA extraction control, and mock community, on each plate.

The prepared plates of the Test Phase DNA samples were submitted to Génome Québec (Montréal, Québec) for ITS amplicon generation and sequencing (Bell *et al*., 2016; Lay *et al*., 2018). In order to preferentially target the oomycete community from other eukaryotes and ensure sufficient quantity for detection, a semi-nested PCR strategy was used to generate the ITS1 amplicons from each Test Phase DNA sample (Sapkota & Nicolaisen, 2015). Each sample was used as template in a PCR reaction consisting of 15 cycles using the ITS6 and ITS4 primers (Cooke *et al*., 2000; Sapkota & Nicolaisen, 2015) to amplify a ~1 kbp fragment. A second round PCR reaction using the ITS6 and ITS7ae primers (Sapkota & Nicolaisen, 2015; Taheri *et al*., 2017) was then done to amplify a ~350 bp fragment of the ITS1 region. These amplicons were then prepared for paired-end 250 bp sequencing using Illumina’s MiSeq platform and the MiSeq Reagent Kits v3 (600-cycles) (Génome Québec, Montréal) (Bell *et al*., 2016; Lay *et al*., 2018). We estimated this should provide a mean of 40 000 reads per sample, which is in line with previous studies that describe microbial eukaryote communities (Bell *et al*., 2016; Lay *et al*., 2018; Blakney *et al*., 2022).

### 2.6 Estimating ASVs from MiSeq ITS amplicons

The ITS amplicons generated by Illumina MiSeq sequencing were used to estimate the diversity and structure of the oomycete communities present in both the rhizosphere and roots of each Test Phase *Brassicaceae* sample. The integrity and totality of the ITS MiSeq data downloaded from Génome Québec, all 17 656 076 reads, was confirmed using their MD5 checksum protocol (Roy *et al*., 2018). Subsequently, all data was managed, and analyzed in R (4.0.3 R Core Team, 2020), and plotted using `ggplot2` (Wickham, 2016).

Instead of generating OTUs from the ITS amplicon data, we opted to use DADA2 for ASV inference, as it generates fewer false-positives than OTUs, reveals more low-abundant, or cryptic, microbes, and as ASVs are unique sequence identifiers, they are directly comparable between studies, unlike OTUs (Callahan *et al*., 2016a & 2017; Fitzpatrick *et al*., 2018). First, due to the variable length of the ITS region, `cutadapt` (Martin, 2011) was used to carefully remove primer sequences from all the ITS reads generated from the control samples, the mock communities, and the experimental Test Phase *Brassicaceae* samples, including any primer sequences generated due to read-through. The `filterAndTrim` function from the `dada2` package (Callahan *et al*., 2016a) was then used for all reads, following the default settings, including, removing reads shorter than 50 nucleotides, or of low-quality (Q = < 20). Filtered and trimmed reads were then processed through DADA2 for ASV inference (Fig. S2 & S5). Default settings were used throughout the DADA2 pipeline, except the DADA inference functions `dadaF` and `dadaR` which used the `pool =’pseudo’` argument, to increase the likelihood of identifying rare taxa. Consequently, the chimera removal function `removeBimeraDenovo` included the `method =’pooled’` argument (Callahan *et al*., 2016b).

The unique ASVs inferred from the ITS amplicon data were assigned taxonomy, to the species level when possible, using the UNITE database for all eukaryotes (Abarenkov *et al*., 2020), and the quality of the data was assessed using the included controls (Fig. S5B). Any ASVs assigned to the taxa Viridiplantae, Alveolata, Fungi, Heterolobosa, or Metazoa were subsequently removed. Rarefaction curves confirmed that we captured the majority of the oomycete communities in both root and rhizosphere samples from both field trials (Fig. S6). The experimental Test Phase *Brassicaceae* ITS sequencing data was subsequently re-analysed independently following the described protocol to avoid any biases from the four no-template negative controls, and the four mock communities. These are the Test Phase oomycete ASVs which are reported hereafter.

### 2.7 Inferring phylogenetic trees

We assembled phylogenies for each compartment, from both trials, in order to infer phylogenetic diversity of the Test Phase *Brassicaceae* oomycete communities. Following the method described by Callahan *et al*., 2016b, the ITS region sequences for each ASV inferred from the Test Phase *Brassicaceae* data were aligned using a profile-to-profile algorithm (Wang & Dunbrack, 2004) with a dendrogram guide tree using the `decipher` package (Wright, 2016). With the `phangorn` package (Schliep, 2011), the maximum likelihood of each site was calculated using the `dist.ml` function using a JC69 equal base frequency model, before assembling phylogenies using the neighbour-joining method. An optimized general time reversible (GTR) nucleotide substitution model was fitted to the phylogeny using the `optim.pml` function. Phylogenies were subsequently added to each `phyloseq` object.

### 2.8 α-diversity of the Test Phase *Brassicaceae* rhizosphere and root communities

First, to visualise taxonomic diversity, ASVs were plotted as taxa cluster maps using `heat_tree` from the `metacoder` package (Foster *et al*., 2017) for the rhizosphere and roots of both experiments, where nodes represent class to genus: node colours represent the number of unique taxa, while node size indicates the relative abundance of each ASV. Taxa cluster maps facilitate visualizing abundance, as well as diversity across taxonomic hierarchies (Foster *et al*., 2017).

Second, in order to estimate the coverage of the oomycete class, we incorporated the oomycete phylogenies into the `phyloseq` object following the method described by Callahan *et al*., 2016b. Faith’s phylogenetic diversity was calculated as an α-diversity index from the Test Phase *Brassicaceae* samples using the `pd` function from the `picante` package (Kembel *et al*., 2010; sum of all branch lengths separating taxa in a community). For comparison, Simpson and Shannon’s α-diversity indices were also calculated (Fig. S7).

We assessed differences between the mean phylogenetic diversity between soil histories, and *Brassicaceae* hosts using the non-parametric Kruskal-Wallis rank sum test, `kruskal.test`, as the transformed data did not respect the assumptions for normality. Specific groups of statistical significance were identified with the post-hoc pairwise Wilcoxon Rank Sum Tests, `pairwise.wilcox.test`, with the FDR correction on the p-values to account for multiple comparisons.

### 2.9 Identification of differentially abundant ASVs and specific indicator species

To refine our understanding of the abundance and composition of the Test Phase *Brassicaceae* oomycete communities, we used two complementary methods to identify taxa specific to soil histories, or *Brassicaceae* hosts. First, taxa cluster maps were used to calculate the differential abundance of ASVs between experimental groups, including rhizosphere and root compartments, *Brassicaceae* host plants, and soil histories. Taxa cluster maps were generated using `compare_groups`, in the metacoder package (Foster *et al*., 2017), where the non-parametric Wilcoxon Rank Sum Tests determined if a randomly selected abundance from one group was greater on average than a randomly selected abundance from another group. As the statistical test was performed for each taxon, we used a false discovery rate (FDR) correction on the p-values to account for the multiple comparisons. When the comparison was between more than two groups, the differential abundances were plotted onto the taxa cluster map using `heat_tree_matrix` (Foster *et al*., 2017).

Second, indicator species analysis was used to detect ASVs that were preferentially abundant in pre-defined environmental groups (roots, or rhizosphere, soil histories, or *Brassicaceae* host). A significant indicator value is obtained if an ASV has a large mean abundance within a group, compared to another group (specificity), and has a presence in most samples of that group (fidelity) (Legendre & Legendre, 2012). The fidelity component complements the differential abundance approach between taxa clusters, which only considers abundance. We performed an indicator species analysis for the ASVs identified in the Test Phase of Trial 1, and then Trial 2. From the `indicspecies` package (De Cáceres & Legendre, 2009), we used the `multipatt` function with 9999 permutations. As the statistical test is performed for each ASV, we used the FDR correction on the p-values to account for multiple comparisons.

### 2.10 β-diversity of the Test Phase *Brassicaceae* rhizosphere and root communities

To test for significant community differences between both trials, compartments, soil histories and *Brassicaceae* hosts, we used the non-parametric permutational multivariate ANOVA (PERMANOVA), where any variation in the ordinated data distance matrix is divided among all the pairs of specified experimental factors. The PERMANOVA was calculated using the `adonis` function in the `vegan` package (Oksanen *et al*., 2020), with 9999 permutations, and the experimental blocks were included as “strata”. Our PERMANOVA used a distance matrix calculated with the Bray-Curtis formula and tested the significance of the effects of soil history, *Brassicaceae* host, and compartment.

We used a variance partition, as a complement to the PERMANOVA, to model the explanatory power of soil history, *Brassicaceae* host, and soil chemistry in the structure of the Test Phase *Brassicaceae* oomycete communities. We then quantified how each significant factor (ie, the explanatory variables) impacted oomycete community structure with a distance-based redundancy analysis (db-RDA) (Legendre & Legendre, 2012). First, singleton ASVs were removed before the `phyloseq` data were transformed using Hellinger’s, such that ASVs with high abundances and few zeros are treated equivalently to those with low abundances and many zeros (Legendre & de Cáceres, 2013). With the `vegan` package (Oksanen *et al*., 2020), soil chemistry was standardized (Legendre & Legendre, 2012) using the `decostand` function. We modelled the explanatory power of each experimental factor in each compartment from both experiments with a variance partition of a partial RDA, using the `varpart` function, and a Bray-Curtis distance matrix (Borcard *et al*., 1992). Variation in the oomycete community data not described by the explanatory variables were quantified by the residuals. Finally, to quantify the amount of variation described by each explanatory factor, db-RDA were calculated using the `capscale` function. Colinear variables were only identified in the Trial 2 soil chemistry, such that specific variables were removed without a loss of information. We subsequently removed total carbon from both the Trial 2 root and rhizosphere RDAs, as well as the zinc concentration from the root RDA. The final plots were generated using p`hyloseq` (McMurdie & Holmes, 2013).

### 2.11 Co-inertia analysis of the relationship between oomycetes and bacterial communities

We used a co-inertia analysis (Dolédec & Chessel, 1994; Legendre & Legendre, 2012) to compare how each sample was influenced by different ASVs as a means to evaluate the strength of the soil history effect. Briefly, this analysis identifies the relationship between two datasets from a common sample by projecting that sample into a common multivariate space. For this analysis the two datasets were the oomycete ASVs identified in this study—and were structured by soil history—and the bacterial ASVs identified from the same experiment, and extracted DNA samples, but were not structured by soil history (Blakney *et al*., 2022). This type of analysis is appropriate for exploring relationships in species-rich datasets—for example, where there are more ASVs than sites—and it imposes no assumptions on the datasets, such as cooccurrence, or interactions (Legendre & Legendre (2012).

The analysis identifies the axes of the common co-inertia space that represent the greatest inertia, or spread, of the common data. The analysis then compares how the positions of each sample in the new co-inertia space are influenced by particular bacterial or oomycete ASVs. The direction of the arrows indicates how a sample is influenced by bacterial ASVs (tail) compared to oomycete ASVs (head); samples with shorter arrows are more similar (Legendre & Legendre, 2012; Mamet *et al*., 2017). Co-inertia analysis is also evaluated with a RV co-efficient (R = correlation, V = vectorial); a multidimensional correlation coefficient equivalent to the Pearson correlation coefficient. A higher RV indicates a stronger relationship between the oomycete and bacterial community matrices (Legendre & Legendre, 2012; Iffis *et al*. 2016).

Therefore, if soil history was particularly significant to the relationship between the bacterial and oomycete ASVs, the samples might be clustered by soil history on the common co-inertia space. Alternatively, a weaker soil history may only be reflected in the shift from bacterial ASVs to oomycete ASVs. This might be plotted by longer arrows oriented toward common oomycete ASVs, which represent the different by soil histories. Finally, if soil history has little to do with the relationship between the bacterial and oomycete ASVs, there may be no discernable pattern in how the sample are plotted, and the arrows between communities would be short.

First, to facilitate the analysis we reduced the bacterial p`hyloseq` objects by removing any ASVs that occurred only once. The p`hyloseq` objects for the oomycete and bacterial communities from the roots and rhizosphere from both field trials were transformed using Hellinger’s transformation. Finally, each oomycete-bacterial sample pair were subjected to co-inertia analysis using the `coinertia` function from the `ade4` package (Dray & Dufour, 2007). The large number of ASVs identified here and from Blakney *et al*., 2022, precluded us from plotting the ASVs onto the co-inertia plane. However as noted in Legendre & Legendre (2012) these are not an essential to the co-inertia analysis.

## 3. Results

### 3.1 Identifying the oomycete communities from the *Brassicaceae* roots and rhizosphere

To identify the composition of the oomycete rhizosphere and root communities from the Test Phase *Brassicaceae* crop species, we inferred ASVs from the retained ITS amplicons using the DADA2 pipeline (Callahan *et al*., 2016 & 2017). The four replicates of the oomycete mock community were sequenced to an average of ~23 000 reads, and closely resembled each other in ASV composition (Fig. S5B & C). From the retained MiSeq reads of the mock community, DADA2 inferred 6 individual ASVs which were identified as 2 *Pythiales*, 2 *Peronosporales*, 1 *Saprolegniales*, and 1 *Albuginales* (Fig. S5B). The mock community was composed of 21 individuals from these four orders (Table S3); this provides some reassurance that our pipeline was effective in identifying the oomycetes present in each experimental sample.

We retained 8 222 283 high-quality ITS MiSeq amplicons through the pipeline, with more reads retained in the rhizosphere samples of the Test *Brassicaceae* crop species in both Trials (Table 1; Fig. S5). 1037 ASVs were inferred from the retained reads, which were subsequently filtered to 412 oomycete ASVs identified among the Test Phase samples. Differences between the rhizosphere and root Test Phase oomycete communities were highly significant in both field trials (Trial 1 PERM *R*^2^ = 0.1226, *p* < 0.0001; Trial 2, PERM *R*^2^ = 0.0751, *p* < 0.0001). The majority of oomycete ASVs were found in the rhizosphere compared to their cognate root samples (Table 1). The oomycete rhizosphere communities were also consistently more phylogenetically diverse than the root communities (Fig. 1), which reflects the greater species richness observed in the rhizosphere (Fig. S7).

**Figure 1.**
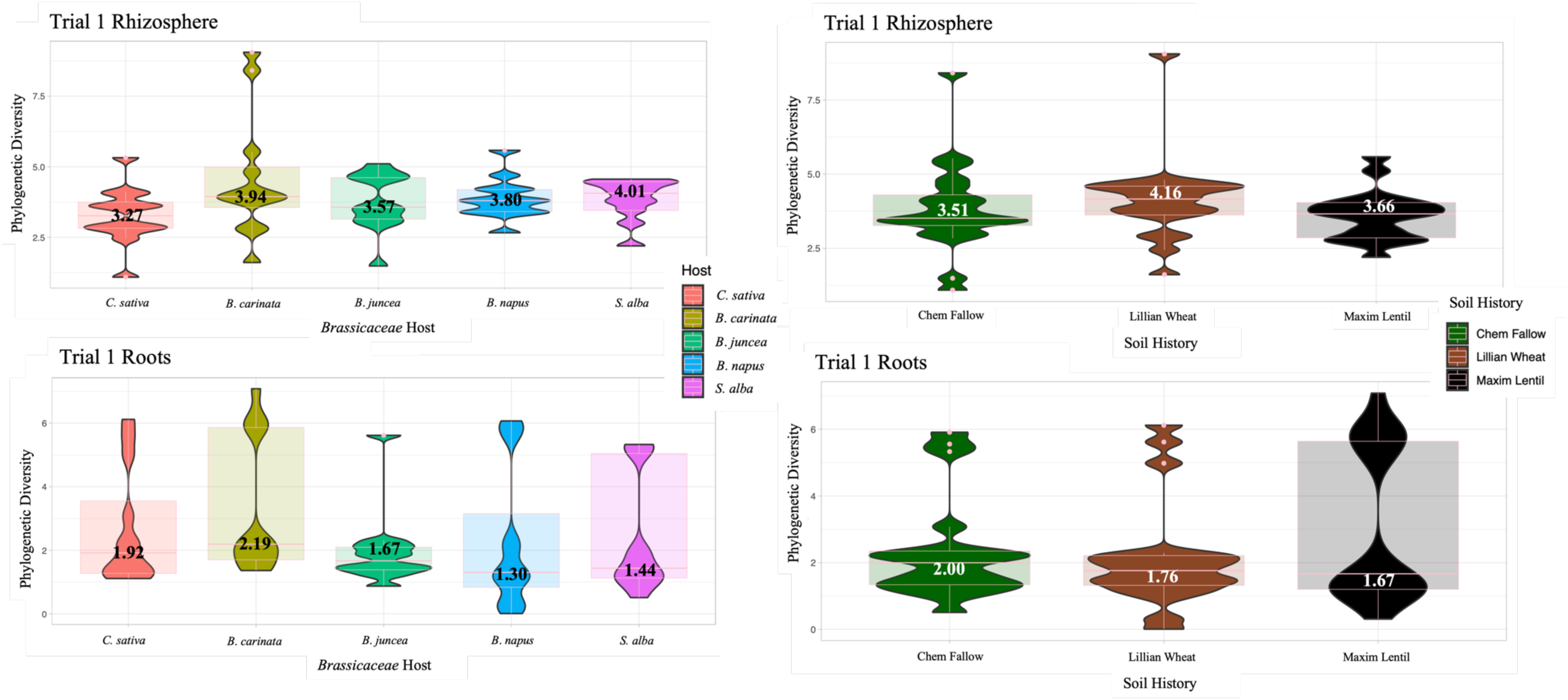
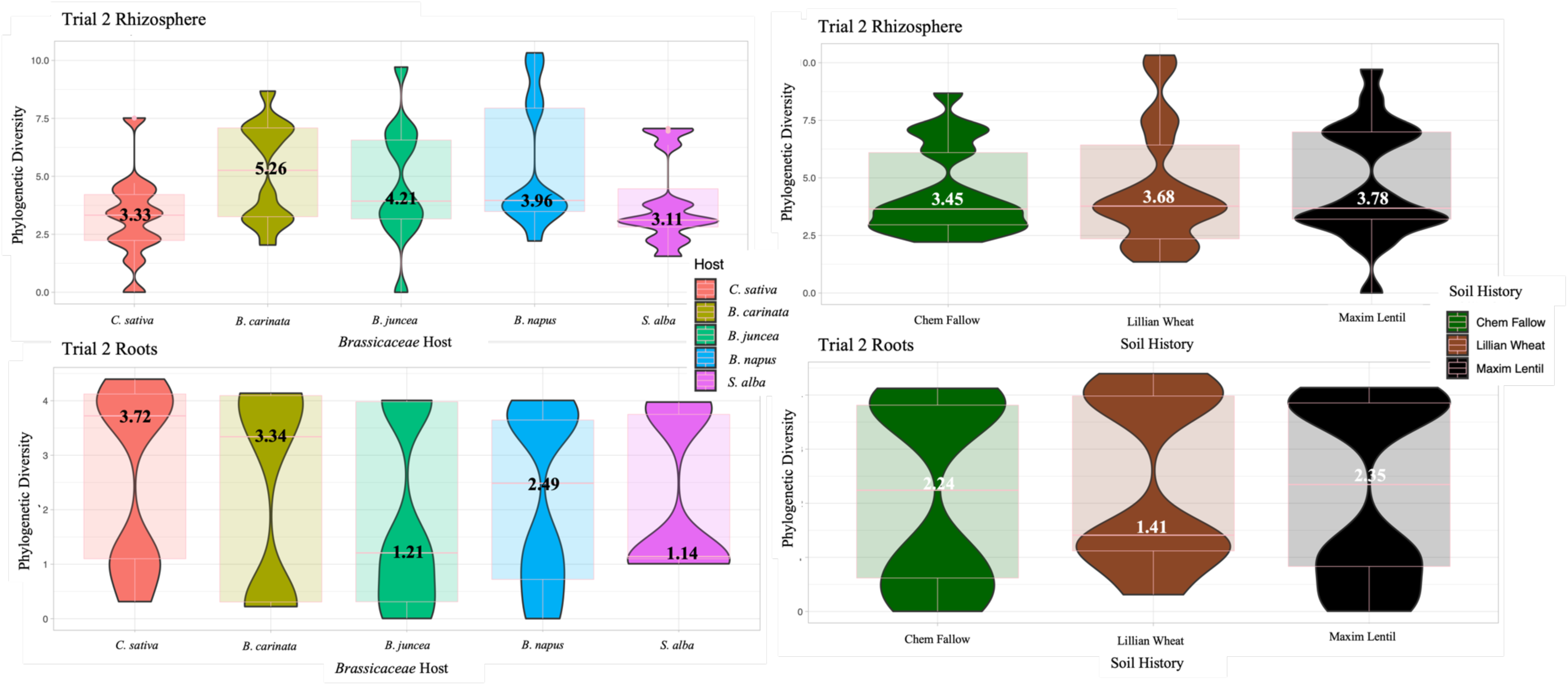
Phylogenetic diversity tended to be higher in the oomycete rhizosphere communities than in the root communities from five *Brassicaceae* host plants in the Test Phase of a two-year rotation, harvested in 2016 (A, Trial 1) and 2017 (B, Trial 2) from Swift Current, Sask. Phylogenetic diversity also tended to be higher in the *Brassicaceae* host plants in Trial 2 than Trial 1. Diversity tended to be higher in the root communities in *Camelina sativa* compared to the corresponding rhizosphere communities only in Trial 2 (B). As the transformed data did not adhere to assumptions of normality, the non-parametric Kruskal test was used to test for significance among the Test Phase oomycete communities grouped by *Brassicaceae* host crops, or by their Conditioning Phase soil histories; no significant differences were detected.

When the oomycete ASVs were plotted as taxa clusters, we observed similar taxonomic composition between the oomycete Test Phase rhizosphere and root communities from Trial 1 and 2 (Fig. 2). *Pythium* species dominated the roots and rhizosphere in both trials in terms of relative abundance, while the order *Peronosporales* consistently had the most taxa across each community (Fig. 2). In Trial 1, *Pythium* and *Peronospora* genera were significantly enriched (*p* < 0.01) in the Test Phase rhizosphere communities compared to the roots, while in Trial 2, only *Pythium* species were significantly more abundant in the rhizosphere (Fig. S8).

**Figure 2.**
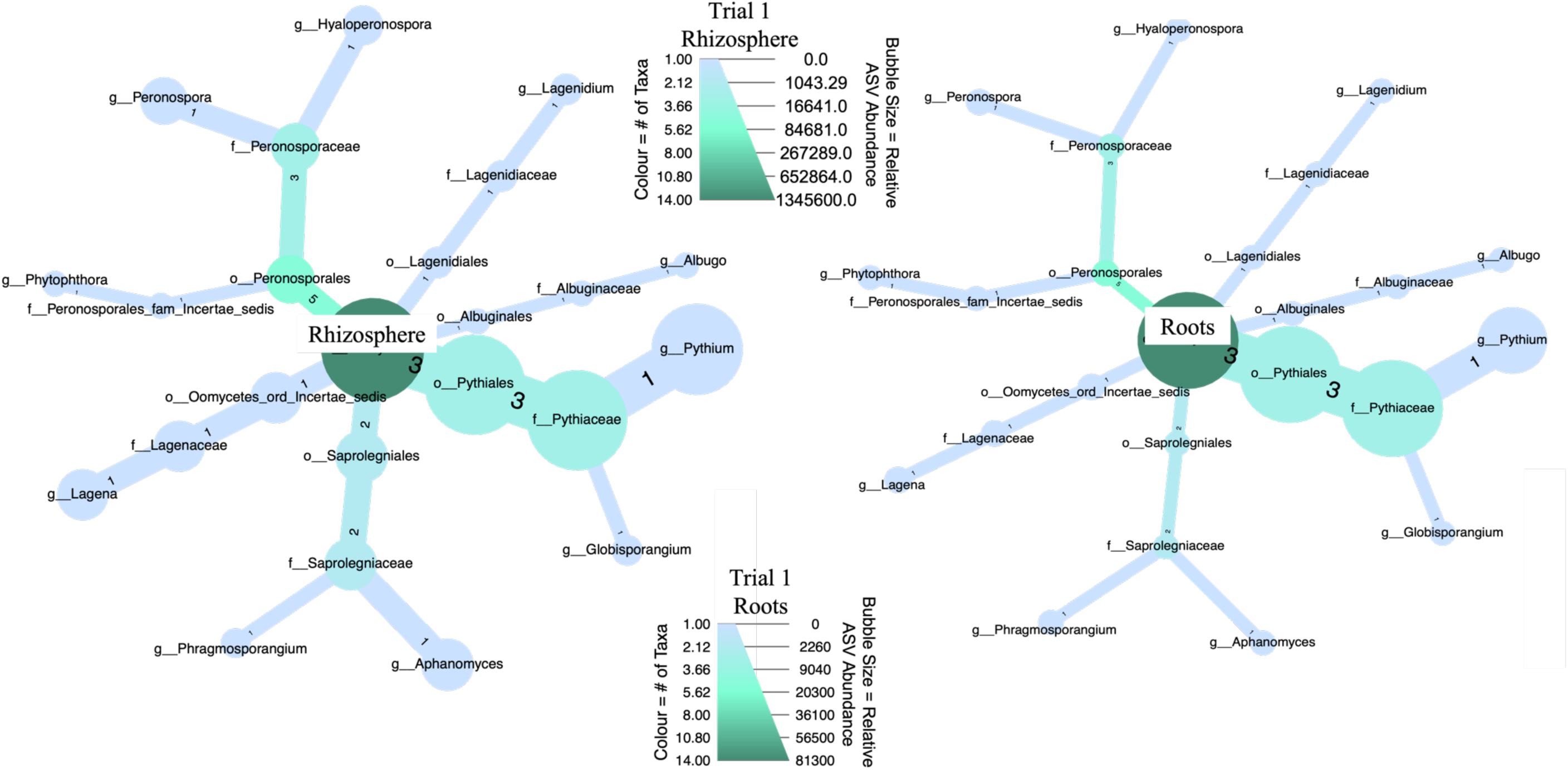
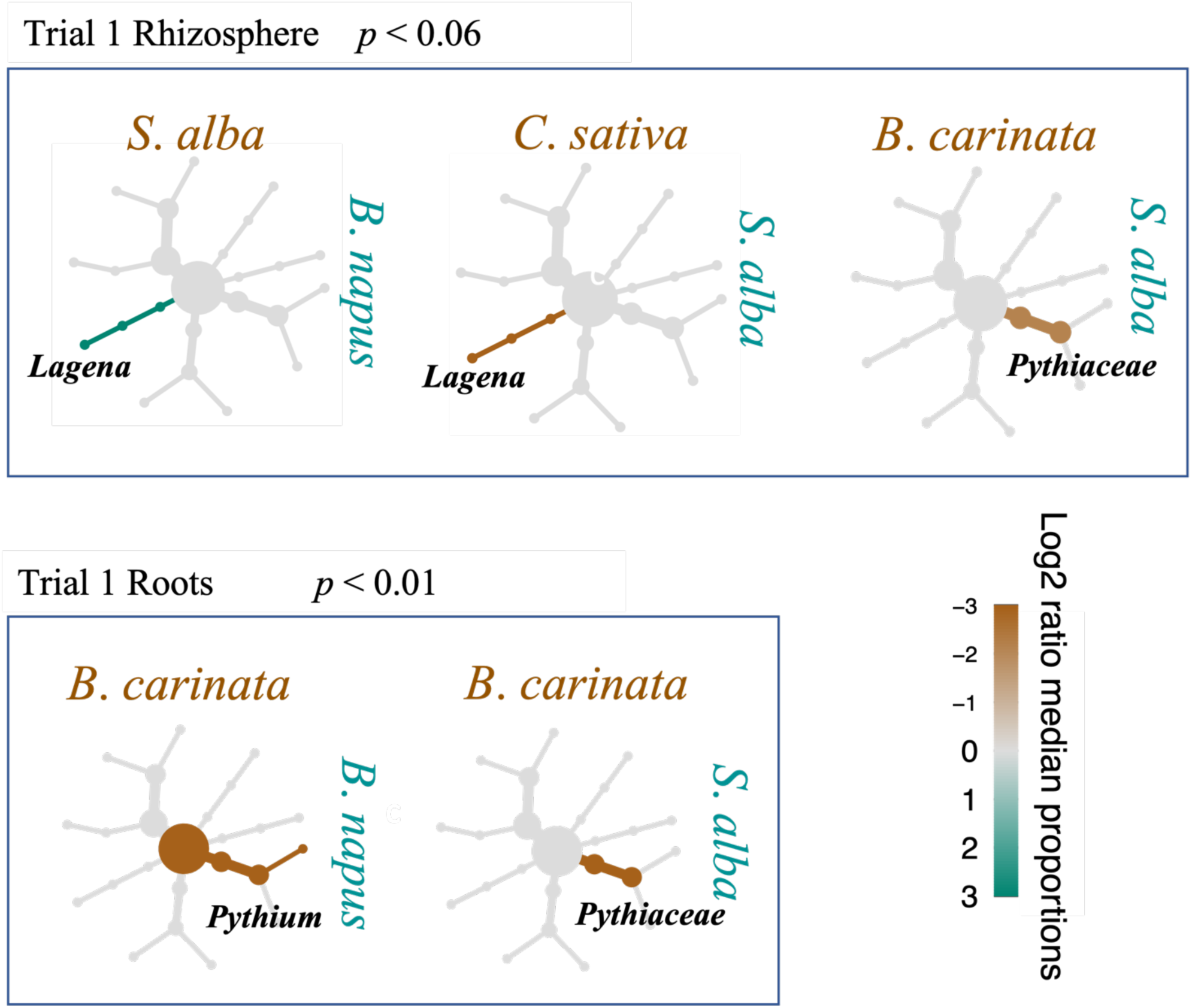
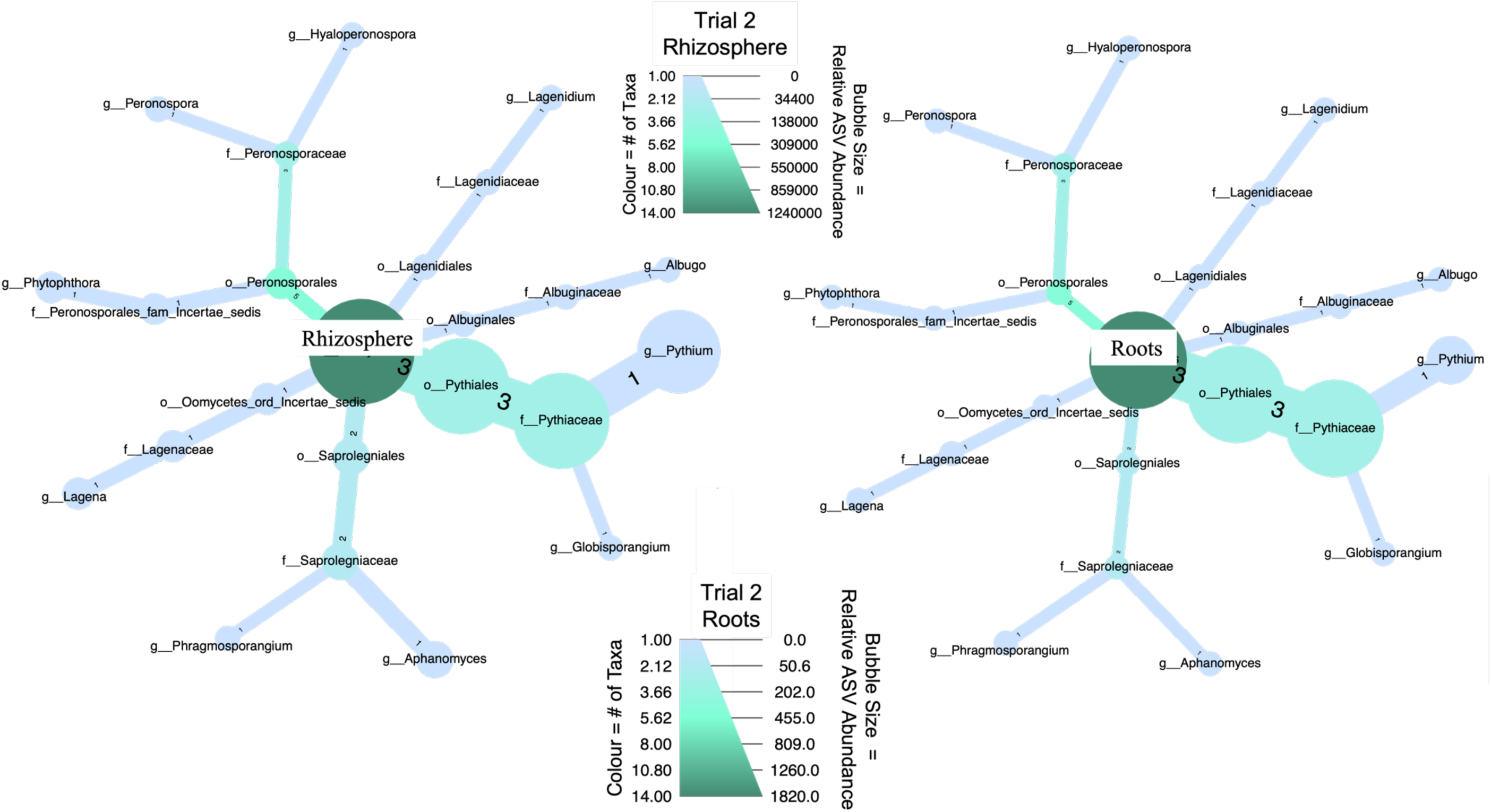
Taxa clusters of the oomycete ASVs inferred from among the rhizosphere and roots of five *Brassicaceae* host crops in the Test Phase of a two-year rotation from Swift Current, SK. The abundance and composition of the oomycete communities are represented to the genus level, where the size of each taxonomic group (bubble) represents the abundance of inferred ASVs, and the colour scale represents the number of unique taxa. (A) In Trial 1 (harvested 2016), *Pythium* dominated in both the rhizosphere (left) and root (right) communities, while the genera *Aphanomyces*, *Lagena*, and *Peronospora*, were dramatically reduced between the rhizosphere and the root communities. (B) Significantly enriched taxa, labelled in bold, were identified between each pair of *Brassicaceae* host crops in Trial 1 rhizosphere (top panel), and root (bottom panel). Taxa that were significantly more abundant are highlighted brown or green, following the labels for each compared factor. Differential taxa clusters identified significantly enriched (ie, abundant) taxa, using the non-parametric Kruskal test, followed by the post-hoc pairwise Wilcox test, with an FDR correction. No enrichment was detected for Trial 2, nor did soil histories enrich any taxa in either experiment. (C) In Trial 2 (harvested 2017), similar to A, *Pythium* dominates in both the rhizosphere and root communities, though there are very few oomycete ASVs detected in the *Brassicaceae* root communities.

Indicator species analysis identified oomycete ASVs specific to the Test Phase rhizosphere communities in both trials, but none in the root communities. Forty-one ASVs were specific to the oomycete rhizosphere communities in Trial 1 (*p* < 0.005, Table 2), while no ASVs were specific to the root communities. Thirty-four ASVs belonged to the *Pythiaceae*, of which half were further identified as *Pythium* sp., two ASVs were *Lagena* sp., and one was *Aphanomyces* sp. (Table 2). The final four indicator ASVs were unknown oomycetes (Table 2). In Trial 2, indicator ASVs in the rhizosphere communities were similar to those from the rhizosphere of Trial 1; 45 ASVs were specific to the Test Phase rhizosphere communities in Trial 2 (*p* < 0.05, Table 2), whereas none were identified in the cognate root communities. The *Pythiaceae* accounted for 37 of these ASVs, of which 19 were further identified as *Pythium* sp (Table 2). Two ASVs were recognized as *Lagena* sp., one was *Aphanomyces* sp., while the remaining five ASVs were unknown oomycetes (Table 2).

### 3.2 The soil history effect on oomycete communities from the roots and rhizosphere of *Brassicaceae* crops

Next, we tested if the three soil histories established by the previous crops structured significantly different oomycete communities. Soil history (Trial 1, PERM *R^2^*= 0.0292, *p* < 0.0015; Trial 2, PERM R^2^ = 0.0300, *p* < 0.0018) was significant in structuring the Test Phase oomycete communities in both field trials (Table 3), though the soil history ~ crop host interaction was not significant in either Trial. We complemented the PERMANOVA by a variance partition to model the explanatory power of each factor (soil histories, soil chemistry, and the *Brassicaceae* host crops). Variance partitioning found that soil history explained similar amounts of the oomycete rhizosphere community data in both field trials (Trial 1 *R^2^* = 0.0453, *p* < 0.001, Fig. 3A; Trial 2 *R^2^* = 0.0476, *p* < 0.001, Fig. 3C). We also quantified how the experimental factors impacted the oomycete community structure with an RDA. Soil history was highly significant for the Test Phase oomycete rhizosphere communities in both field trails (Trial 1 adj. *R*^2^ = 0.0539, *p* < 0.001, Fig. 4A; Trial 2 adj. *R^2^*= 0.0727, *p* < 0.001, Fig. 4B), where the communities were grouped by soil history (Fig. 4).

**Figure 3.**
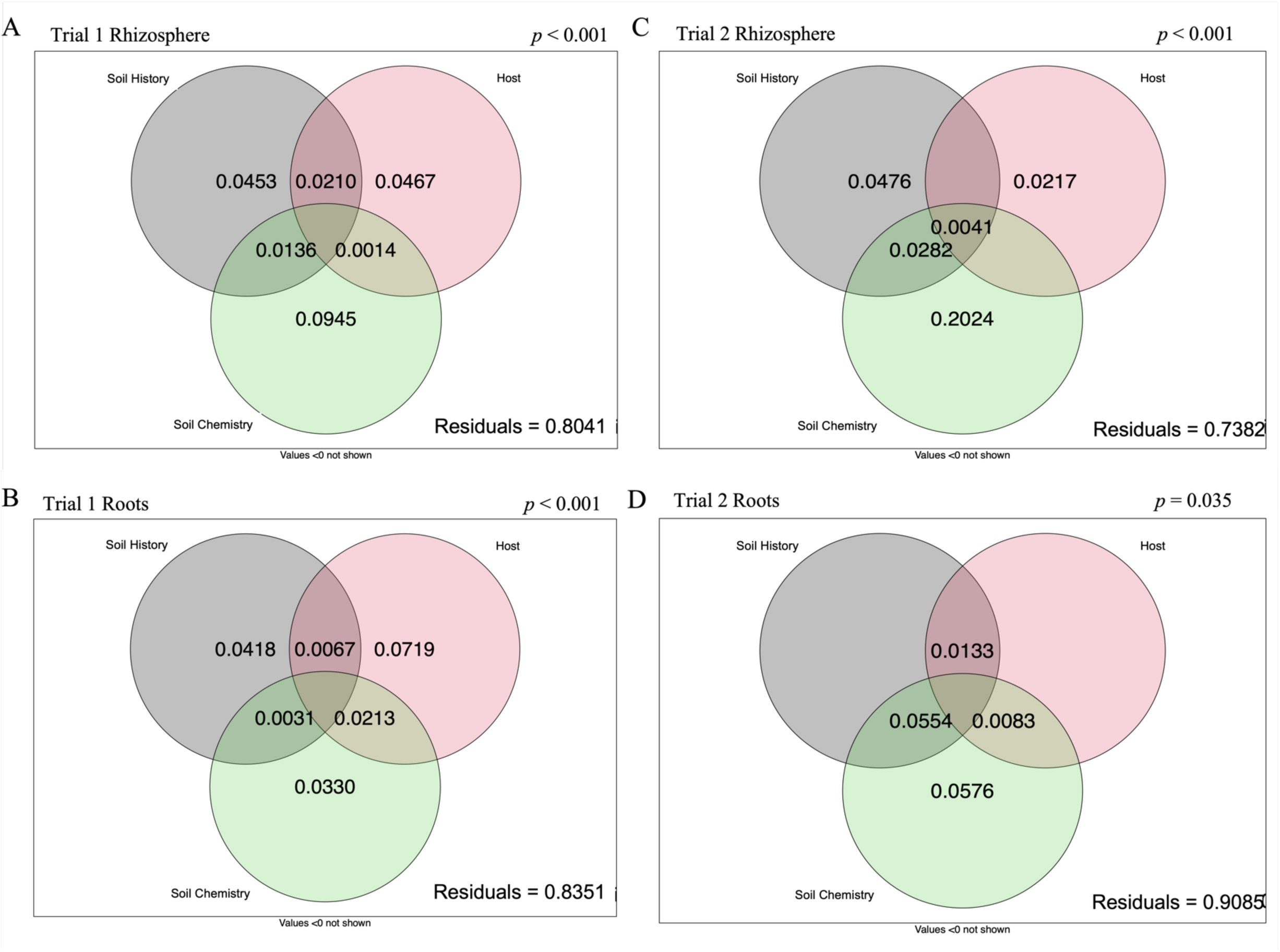
Soil chemistry, soil history, and the *Brassicaceae* host crops were each influential in structuring the oomycetes communities in the rhizosphere and roots, from the Test Phase of a two-year crop rotation, harvested in 2016 (Trial 1) and 2017 (Trial 2) from Swift Current, SK. Soil history explained a consistent amount of variance in the oomycete communities (A, B, & C). In the rhizosphere (A & C), the current soil chemistry consistently explained the most variance in the oomycete communities. In Trial 1 (A & B), the influence of the host plants increases over the communities moving from rhizosphere (A) into the roots (B). Bray-Curtis distances were used in the variance partition.

**Figure 4.**
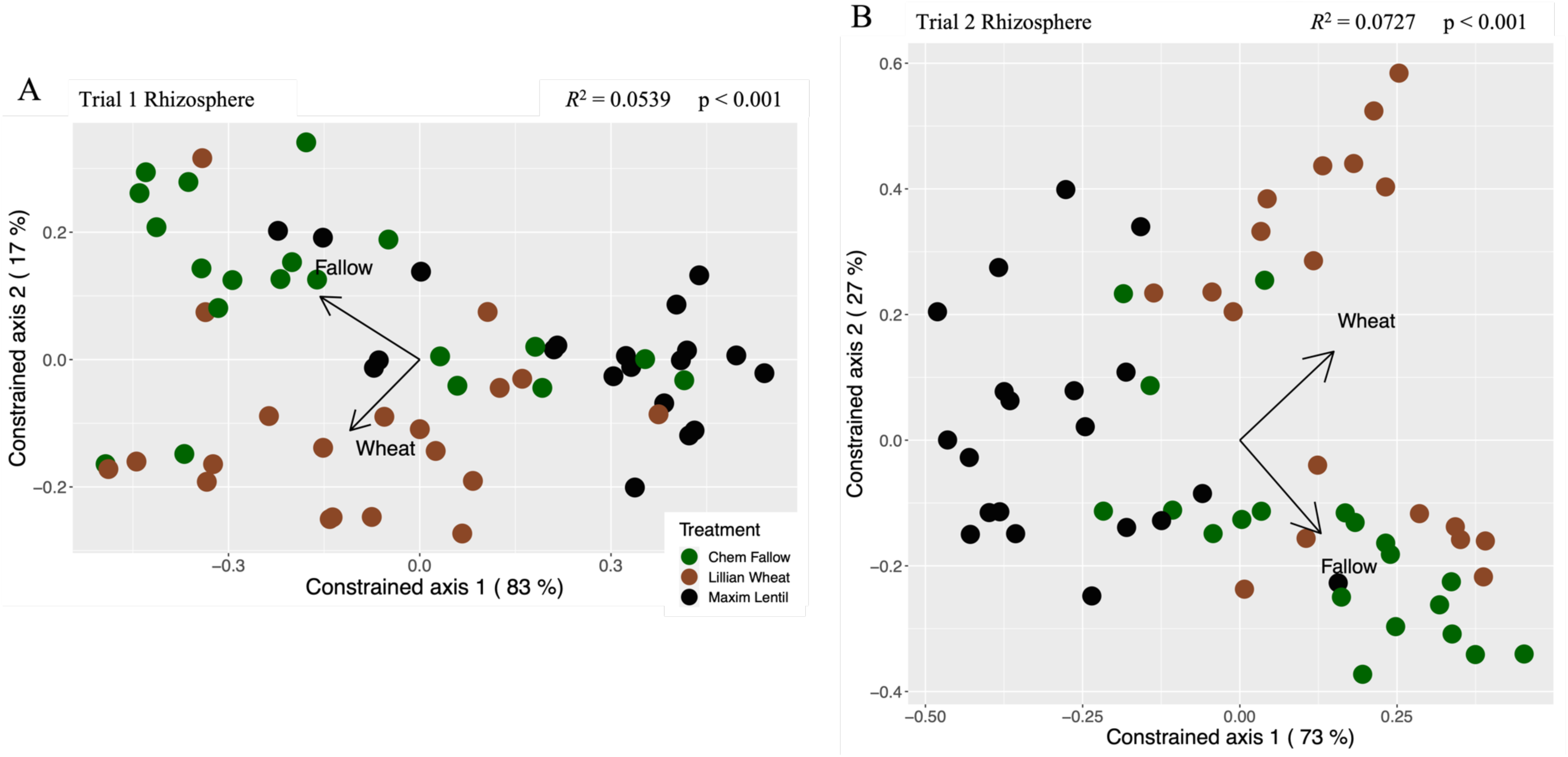
Soil history was significant in structuring the oomycete rhizosphere communities in both field trials of a two-year crop rotation, harvested in 2016 (Trial 1) and 2017 (Trial 2) from Swift Current, SK. Distance-based redundancy analyses quantified how soil history impacted the oomycete community structure, where communities with similar composition appear closer together

Soil history was less consistent in the oomycete root communities of both field trials. First, it explained a similar amount of the variance in the Test Phase root community data (*R^2^* = 0.0418, *p* < 0.001, Fig. 3B) as in the rhizosphere communities in Trial 1, but was not significant in the root community data of Trial 2 (Fig. 3D). Second, RDAs demonstrated the importance of soil history in the oomycete root communities of both trials (Trial 1 adj. *R*^2^ = 0.0429, *p* = 0.007, Fig. S9A; Trial 2 adj. *R^2^*= 0.0403, *p* = 0.005, Fig. S9B). Though they were less significant compared to the rhizosphere (Fig. 4), they explained a similar amount of the data (*R*^2^ = ~0.04, Fig. 4 & S9). A gradient separating the oomycete root communities based on their previous soil history was observed in Trial 2 (Fig. S9B), similar to the corresponding rhizosphere communities (Fig. 4), but was less obvious in Trial 1 (Fig. S9A).

Soil history also determined indicator species identified in the Test Phase rhizosphere communities in both Trials. In Trial 1, five oomycete ASVs were specific to rhizosphere communities with lentil crop histories; four were recognized as *Pythiaceae* and the fifth as a *Pythium* sp. (Table 2). In Trial 2, 11 *Pythiaceae* ASVs were specific to rhizosphere communities with the lentil soil history (*p* < 0.05, Table 2), while another *Pythiaceae* ASV was specific to rhizosphere communities with the wheat soil history (Table 2).

### 3.3 The influence of the *Brassicaceae* crops on their oomycete communities

*Brassicaceae* hosts (*R^2^* = 0.0520, *p* < 0.0003) had a significant effect on oomycete community structure in Trial 1 (Table 3), but not in the dry year of Trial 2. The variance partition illustrated that the *Brassicaceae* crop hosts accounted for 4.67% of variance of the Test Phase rhizosphere community data (Fig. 3A) in Trial 1. However, *Brassicaceae* crop hosts were not significant in the variance partition of Trial 2 (Fig. 3C & D). RDA also supported the significance of the *Brassicaceae* crop host (*R*^2^ = 0.0454, *p* = 0.006, Fig. S10A) in the oomycete rhizosphere communities in Trial 1, but was not significant in Trial 2. The first RDA axis showed a gradient among oomycete communities between *C*. *sativa* and *S*. *alba*, with a notable amount of overlap. Interestingly, the second axis showed a gradient between soil histories, with the majority of communities from lentil sites clustered in the bottom left (Fig. S10A).

In the oomycete root communities, *Brassicaceae* crop hosts explained the most variation in Trial 1 (*R*^2^ = 0.0719, *p* < 0.001, Fig. 3B), but were not significant in the variance partition of the root communities in Trial 2, similar to the rhizosphere in Trial 2. RDA also illustrated the importance of the *Brassicaceae* crop hosts in structuring the Test Phase oomycete root communities (*R*^2^ = 0.0961, *p* = 0.002, Fig. S10B) in Trial 1, but not in Trial 2. Oomycete root communities from *C*. *sativa* and *B*. *carinata* had more distinct clusters than the other three crop hosts in Trial 1 (Fig. S10A).

Differential taxa clusters from Trial 1 identified variations between *Brassicaceae* hosts in Trial 1, but not in the dry year of Trial 2. The Test Phase oomycete communities in *S*. *alba* rhizospheres were depleted in *Lagena* sp. ASVs relative to the rhizosphere communities of both *B*. *napus* and *C*. *sativa* (*p* < 0.06, Fig. 2B). *S*. *alba* root (*p* < 0.01) and rhizosphere (*p* < 0.06) communities were also depleted in *Pythiaceae* ASVs relative to the oomycete communities of *B*. *carinata* (Fig. 2B). Test Phase oomycete root communities from *B*. *carinata* were enriched in *Pythium* sp. ASVs, compared to *B*. *napus* (*p* < 0.01, Fig. 2B). Indicator species analysis did not identify any oomycete ASVs as specific to any of the five *Brassicaceae* host crops in either field trial.

### 3.4 Soil chemistry on the oomycete rhizosphere and root communities

Variance partitioning revealed that the soil chemistry was the most significant factor in the oomycete rhizosphere communities in both field trials (Trial 1 *R^2^* = 0.0945%, *p* < 0.001, Fig. 3A; Trial 2 *R^2^* = 0.2024, *p* < 0.001, Fig. 3C). RDA also supported that soil chemistry was the most explicative experimental factor of the Test Phase oomycete rhizosphere communities in both trials (Trial 1 adj. *R*^2^ = 0.1004, *p* = 0.012, Fig. 5A; Trial 2 adj. *R^2^* = 0.271, *p* < 0.001, Fig. 5B). These data indicate that the Test Phase oomycete rhizosphere communities were strongly shaped by soil chemistry in both field trials. This effect was stronger in the rhizosphere during the dry year of Trial 2.

**Figure 5.**
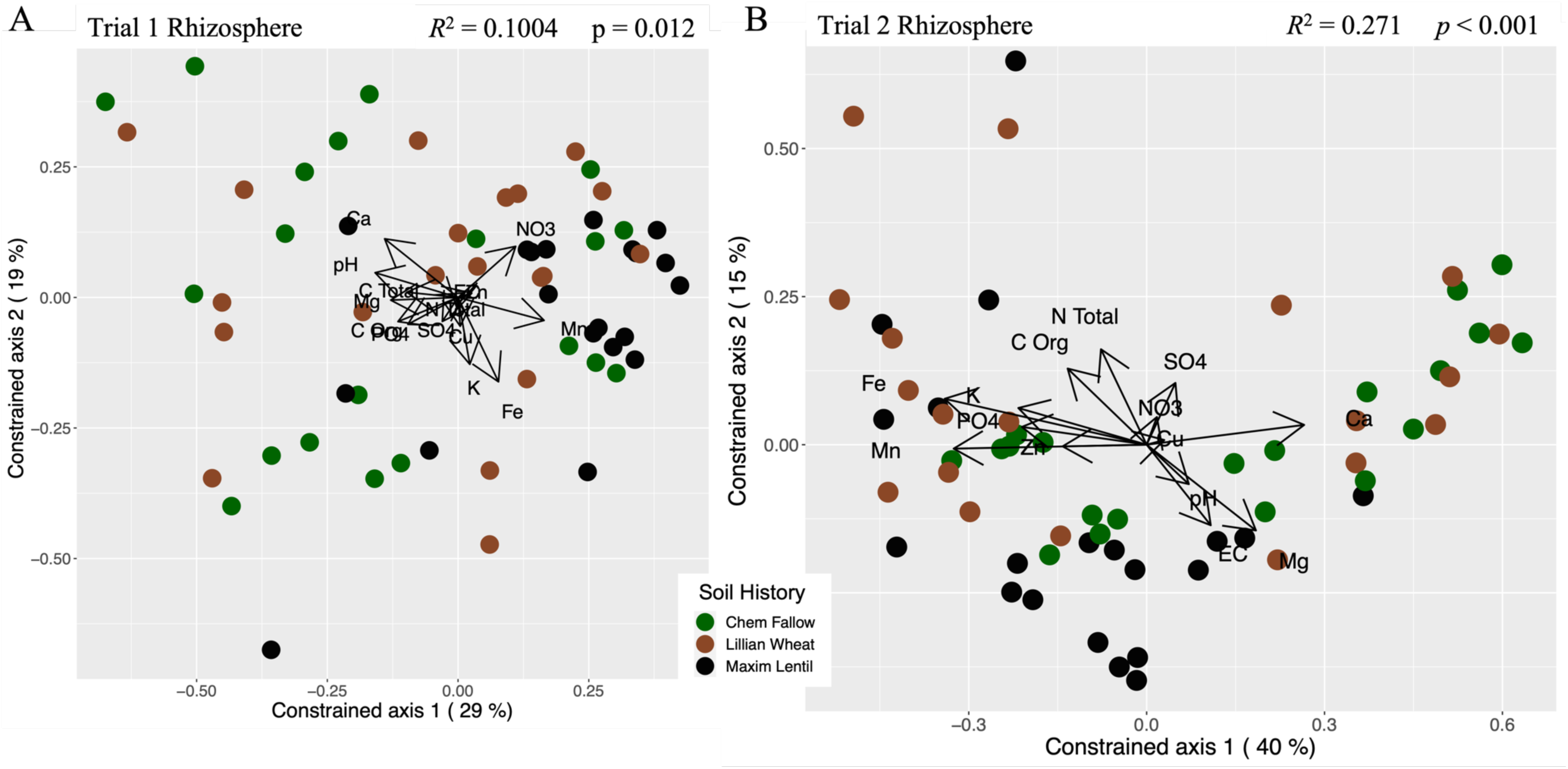
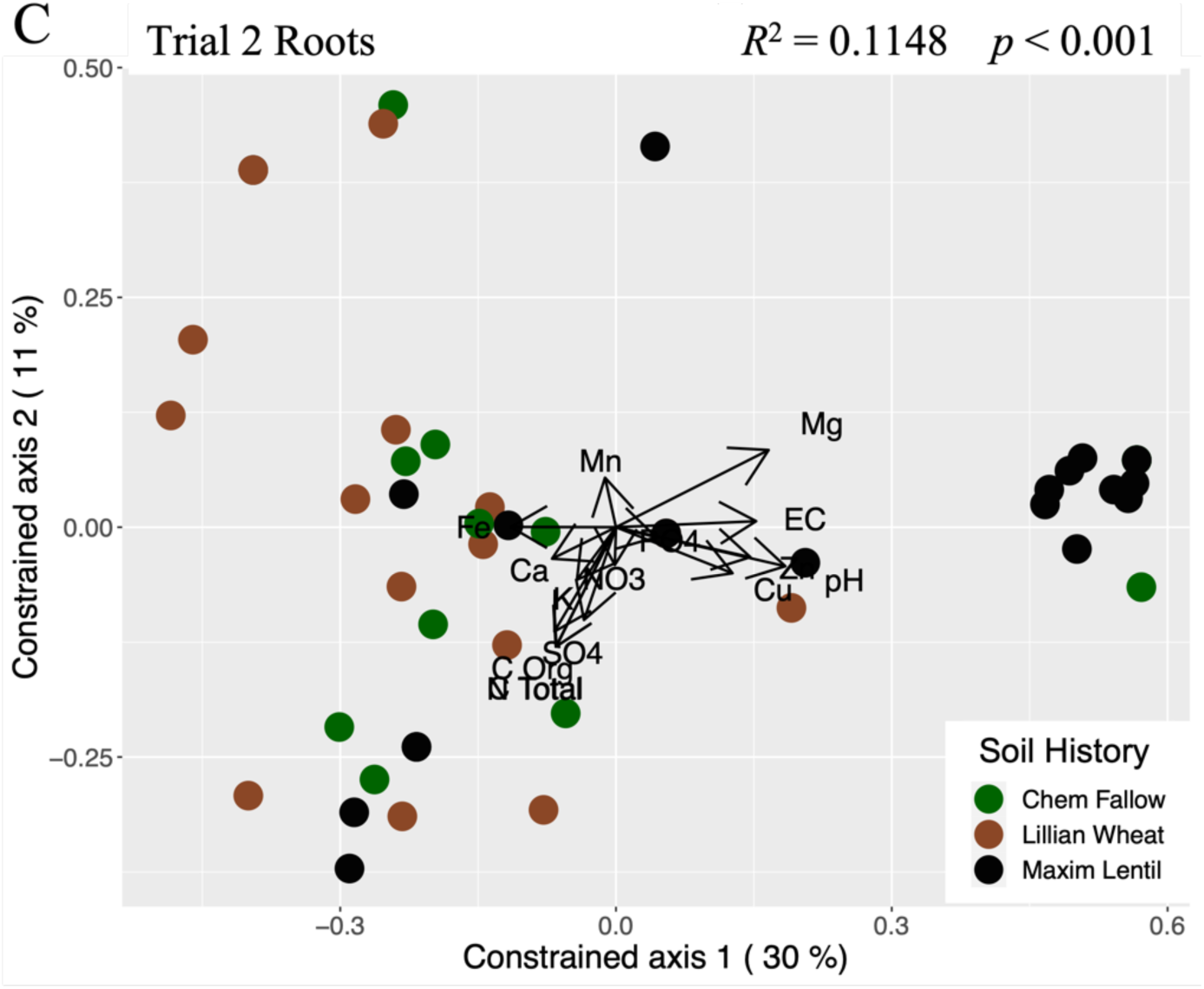
Soil chemistry was the most significant factor structuring oomycete community structures in the rhizosphere (A, *R^2^*= 0.1004, *p* − 0.012; B, *R^2^* = 0.271, *p* < 0.001) from both field trials, as well as in the roots from five *Brassicaceae* crop hosts from Trial 2 (C, *R^2^* = 0.1148, *p* < 0.001) in the Test Phase of a two-year crop rotation, harvested in 2016 (Trial 1), and 2017 (Trial 2) from Swift Current, SK. Distance-based redundancy analyses quantified how soil chemistry structured oomycete community structure, where communities with similar composition appear closer together. The largest factors in the rhizosphere were (A): iron, calcium, nitrate, and manganese, which contrasted with pH; and (B) calcium contrasting with manganese, and to a smaller extent zinc, magnesium opposed organic carbon, conductivity and pH contrasted total nitrogen; iron was also a strong factor. (C) the largest factor in the root communities were magnesium, which was weakly contrasted with calcium, while conductivity contrasted with iron. Note that in B and C total carbon was explained by organic carbon, while Zn was explained by Cu in C.

In the oomycete root communities, soil chemistry explained the least amount of variation in Trial 1 (*R^2^* = 0.0330, *p* < 0.001, Fig. 3B), but was the only significant factor during the dry year in Trial 2 (*R^2^* = 0.0576, *p* < 0.035, Fig. 3D). RDA also supported the significance of soil chemistry on the Test Phase oomycete root communities in Trial 2 (adj. *R^2^* = 0.1148, *p* < 0.001, Fig. 5C), but not in Trial 1. The data suggests that soil chemistry was only influential in the oomycete root communities during the dry year of Trial 2, when the effects of soil history and *Brassicaceae* crop host were reduced on structuring the communities.

### 3.5 Co-inertia analysis between the *Brassicaceae* oomycete and bacterial communities

We used a co-inertia test to investigate how influential soil history was on the relationship between the Test Phase oomycete communities investigated here and the previously identified bacterial communities (Blakney *et al*., 2022). In Trial 1, the root and rhizosphere samples had similar RV coefficients, 0.6291 and 0.7049, respectively, suggesting that the oomycete and bacterial communities did have significant relationships in both compartments. The rhizosphere samples were plotted on the first and second axes which represented 8.185% and 6.551% of the co-inertia. The low inertia suggests little influence of the two sets of ASVs on the samples. This is further illustrated in that the majority of the 60 samples in Trial 1 appeared quite similar, as they remained clustered together toward the centre of the plot. Rhizosphere samples from plots 17, 41, 46, and 51, were relatively more influenced by the presence of particular microbial ASVs, as they are further from the centre (Fig. S11A). Only samples 17 and 51 illustrated any noticeable divergence between which microbial ASVs influenced their structure, given their appreciable arrow lengths.

The root samples from Trial 1 captured 9.404% and 7.682% of the co-inertia in the first and second axes, respectively, which suggests minimal influence of the ASVs on each sample, similar to their corresponding rhizosphere communities. Most of the root samples were also clustered together in the co-inertia plot; only samples 47 and 51 were noticeably influenced by the presence of particular microbial ASVs, given their positions. The arrow length of the root sample from plot 47 suggests it diverged between which microbial ASVs influenced their structure (Fig. S11B).

In Trial 2, the root and rhizosphere samples had less similar RV coefficients; the oomycete and bacterial communities had a significant relationship (RV = 0.8307) in the rhizosphere samples, but not in the root samples (RV = 0.5767). The first and second axes for the rhizosphere samples represented 12.776% and 4.578% of the co-inertia, respectively. Given the low inertia the rhizosphere samples appeared to be weakly influenced by the two datasets of microbial ASVs. Nonetheless, Trial 2 rhizosphere samples were more heterogenous as they were more dispersed compared to the rhizosphere samples from Trial 1, with samples 22 and 30 being distinctly different (Fig. S11C). Finally, the influence of the microbial ASVs only appeared to shift in samples 19 and 30, as shown by their arrow lengths.

The Trial 2 root samples captured 8.880% and 8.131% of the co-inertia in the first and second axes, respectively. Again, the low inertia suggests minimal influence of microbial ASVs on each sample, which is similar to their corresponding rhizosphere communities. Unlike the rhizosphere samples, however, the root samples were tightly clustered; only sample 51 was noticeably influenced by particular microbial ASVs. The arrow length of sample 51 also suggests the influence of the microbial ASVs shifted between the bacterial and oomycete communities (Fig. S10C).

## 4. Discussion

How the soil history established by previous plant-soil microbial communities’ condition future generations of oomycete communities remains relatively unknown. Oomycetes are vastly understudied compared to bacteria and fungi, yet are important microbial communities, especially for agriculture where many oomycetes are responsible for severe declines in yields. In this study, we investigated the impact of three different soil histories established by the previous year’s crops on the soil oomycete communities associated with five *Brassicaceae* oilseed host crops. The nested-ITS amplicon strategy we incorporated with MiSeq metabarcoding specifically targeted oomycetes, and illustrated that soil history had a greater influence on the communities than the *Brassicaceae* host crops, while soil chemistry structured the oomycete communities more during the dry field trial. Our results highlight the impact of edaphic factors over different growing seasons and the importance of monitoring and quantifying oomycete biodiversity.

### 4.1 Previous crops significantly impacted the oomycete rhizosphere communities

The previous crops, and their agricultural treatments, impacted the subsequent oomycete communities through plant-soil microbial community feedback. We hypothesized that the three soil histories established by the previous crops would structure significantly different oomycete communities, regardless of their current *Brassicaceae* host, in both the roots and rhizosphere. Our data illustrated that this was largely sustained; we found consistent support for soil history influencing the structure of the oomycete rhizosphere communities of both field trials, as well as the root communities in Trial 2 (Table 3, Fig. 3). Moreover, gradient analysis (Fig. 4 & S9) highlighted how different oomycete communities tended to cluster according to the soil histories established by the previous crops, especially among the rhizosphere communities. These are exciting results as they raise more questions about oomycete community dynamics and their interactions with different soil histories established through crop rotations.

In this study, we found that the oomycete communities were significantly structured by each of the three previously established soil histories. We observed little effect from the *Brassicaceae* crop hosts to re-structure the oomycete communities according to their current hosts. Crop rotations have previously been shown to quickly adjust subsequent bacterial communities via plant-microbial community feedback mechanisms (Hamel *et al*., 2018; Blakney *et al*., 2022). Important fractions of bacterial rhizosphere communities tend to be fast-growing and have rapid turn-over, which may allow bacterial communities to be more responsive to the dynamic needs of their host plants (Castrillo *et al*., 2017; Hou *et al*., 2021). Thus, plant-microbial community feedback mechanisms from new host plants appears sufficient to quickly erase the soil history established by a previous plant and modify the bacterial communities (Kaisermann *et al*., 2017; Hannula *et al*., 2021; Blakney *et al*., 2022).

Conversely, experimental evidence has suggested that such feedback mechanisms are not sufficient to alter fungal communities (Kaisermann *et al*., 2017; Hannula *et al*., 2021). One suggestion for why the influence of established soil history varies between bacterial and fungal communities has been due to their different growth rates (Semchenko *et al*., 2018; Hannula *et al*., 2021). Compared to the rapidly growing parts of plant bacterial communities, fungal communities remain more stable through time. Fungal growth seems more steady and less influenced by host plant feedback mechanisms, which limits how responsive fungal communities might be to the influence of new crop hosts (Kaisermann *et al*., 2017; Hannula *et al*., 2021). Although oomycetes are not fungi, and have vastly different evolutionary origins (Fawke *et al*., 2015; Kamoun *et al*., 2015; Schwelm *et al*., 2017), our data illustrates a similar trend, where oomycetes, like fungi, remained relatively unaffected by changes in crop hosts possibly due to their growth rate. Complementary to this idea is that oomycete oospores can persist from year to year and are constitutively dormant, such that not all oospores germinate at the same time even under optimal conditions (Martin & Loper, 1999). This could also help account for why oomycete communities may appear less affected by the influence of the new *Brassicaceae* hosts feedback mechanisms (Kaisermann *et al*., 2017; Hannula *et al*., 2021).

In fact, our results illustrate that the soil history established by the previous lentil and wheat crops helped to structure distinct oomycete communities that were still detectable the following year (Fig. 4 & S9, Table 3). The lentil-specific oomycete rhizosphere community we detected in both field trials may be unsurprising since legumes, including lentils, tend to retain more soil moisture compared to other crops, and soil moisture is a key factor for oomycete growth. Recent studies have also suggested that lentils have an increased vulnerability to oomycete outbreaks (Hwang *et al*., 2015; Rojas *et al*., 2017; Karppinen *et al*., 2020). Finally, *Pythiaceae* have previously been reported in Canadian pea fields (Taheri *et al*., 2017), thus detecting a variety of *Pythiaceae* ASV specific to the lentil soil history seems reasonable.

Conversely, the *Aphanomyces* ASV specific to the rhizosphere communities in Trial 2 with wheat soil history (Table 2) is somewhat more unexpected. Most of the interest concerning the *Aphanomyces* focuses on *A*. *cochlioides* and *A*. *euteiches*, which are well described pathogens specific to sugar beets and legumes, respectively (Diéguez-Uribeondo *et al*., 2009). However, there is a divergent lineage that consists of saprotrophs and opportunistic plant pathogens that are not known to maintain specific hosts (Diéguez-Uribeondo *et al*., 2009). A saprotrophic oomycete capable of degrading wheat residues might explain the *Aphanomyces* ASV identified among Trial 2 rhizosphere communities with wheat soil history.

### 4.2 Oomycete communities were not influenced by *Brassicaceae* crops during the drier field trial

Although our initial hypothesis concerning soil history was largely supported, we did nonetheless observe an influence of the *Brassicaceae* crop hosts on the oomycete communities, but only during Trial 1 (Fig. 3, Table 3), and particularly in their roots (Fig. S10). Plant hosts ought to have the most influence to select for microbial communities in their roots, compared to the rhizosphere, or leaf surface (Gavrin *et al*., 2020; Maciá-Vicente *et al*., 2020). While our data illustrates that the *Brassicaceae* host crops were quite significant in the roots during Trial 1 (Fig. 3 & S9), we did not detect any oomycete ASVs as indicator species from any of the five hosts (Table 2). Furthermore, we observed a considerable amount of overlap among the oomycete root communities, notwithstanding the more distinct clusters of communities from *C*. *sativa* and *B*. *carinata* (Fig. S10). Our data may indicate that the influence of the *Brassicaceae* hosts on the oomycete root communities was insufficient to structure more distinct groups of oomycetes. This weaker effect of the *Brassicaceae* hosts could be due to other competing factors, such as the previous soil history, or current soil chemistry. Alternatively, the close genetic relationship of the five *Brassicaceae* crop hosts may have precluded us from identifying more specific oomycete assemblages (Blakney *et al*., 2022).

Moreover, there was no influence of any of the *Brassicaceae* crops on the oomycete communities during Trial 2 that we observed, despite following identical experimental protocols, and the use of the same agricultural management practices and inputs. The disparate observations between the two field trials could be due to the environmental conditions being 6x drier during Trial 2: 55.0 mm of precipitation versus 328.4 mm in Trial 1 (Blakney *et al*., 2022). The *Brassicaceae* host plants appeared to be restricted in growth due to the dry conditions (Fig. S3), which would also constrain their nutrient uptake from the soil, and rhizodeposition (Fitzpatrick *et al*., 2018). Therefore, if the reciprocal plant-soil microbial community feedback was impaired due to the availability of water, it could account for the absent influence of the *Brassicaceae* hosts on the oomycete communities in Trial 2.

The drier conditions of field trial 2 may also have had an impact on the oomycete community itself. Oomycetes prefer high soil moisture for motility, subsequent infection and growth, and the completion of their life cycle (Martin & Loper, 1999; Fawke *et al*., 2015; Martiny *et al*., 2015; Rojas & Huang, 2018). Our data demonstrates a similar community composition between both field trails, though with reductions in sequencing reads and diversity in Trial 2. This could be evidence for how the drier conditions impacted the community. Quantifying community sizes could help determine this in future experiments.

Nonetheless, the impact of the *Brassicaceae* crop hosts remained limited, as the structure of the oomycete communities remained significantly influenced by the previous crops. This could indicate that these specific crops may not be effective as a strategy to limit the accumulation of potentially pathogenic oomycetes in the soil over the short term. Various crop rotations, including those involving *Brassicaceae*, have been shown to help control phytopathogens by restructuring the microbial communities from one season to the next (Etesami & Alikhani, 2016; Yang *et al*., 2021). Such shifts generally occur through plant-soil feedback processes, such as rhizodeposition, or by producing anti-microbial compounds (Krasnow & Hausbeck, 2015; Lebeis *et al*., 2015; Revillini *et al*., 2016; Korenblum *et al*., 2020; Kawasaki *et al*., 2021; Yu *et al*., 2021). For example, *Brassicaceae* crops produce anti-microbial glucosinolates, which have been used to control phytopathogens, including oomycetes (Krasnow & Hausbeck, 2015). However, our data illustrates that the five *Brassicaceae* crops were unable to sufficiently alter the soil history established by the previous crops, given that the oomycete communities remained significantly structured by soil history. This could suggest that these crop rotations may be an insufficient strategy to control oomycete phytopathogens in the short term.

In addition, we observed that *B*. *carinata* crop hosts were significantly enriched in *Pythiaceae* ASVs in their root and rhizosphere communities compared to the other *Brassicaceae* hosts (Fig. 2B). This could suggest that *B*. *carinata* may be more susceptible to oomycete accumulation. Conversely, we also noted that *S*. *alba* hosts were depleted in *Lagena* and *Pythiaceae* ASVs in their rhizosphere communities (Fig. 2B), compared to the other Brassicaceae crop hosts. This might demonstrate increased resistance to accumulating these oomycetes in their rhizosphere. These two examples warrant further study to help evaluate how effective these particular crop rotations may be at limiting oomycete infections.

Further to this, our study points out three recommendations needed to better understand the phytopathogenicity of oomycetes: first, biodiversity monitoring should inventory the oomycete communities established at the end of each growing season and observe which ASVs persisted in a given plot (Derevnina *et al*., 2016; Goméz *et al*., 2021). Second, quantifying the size of each community would help determine if crop rotations actually limit, or reduce, the growth of the oomycete communities. Third, since the impact of a crop may not be observed during the active growing season (Hamel *et al*., 2018), longer field trials with multiple timepoints could help confirm our findings. These additional steps may provide a more nuanced understanding of the dynamics within oomycete communities and help determine the utility of crop rotations as a strategy to limit the accumulation of oomycete phytopathogens in agricultural soil.

### 4.3 Soil chemistry constrained the oomycete community structure

Although we initially sought to test the influence of soil history on structuring oomycete communities, our data revealed that the soil chemistry had the strongest influence in the rhizosphere during both field trials, and among the root communities during the dry season in Trial 2 (Fig. 3 & 5). In our agricultural setting, soil chemistry was largely a synthesis of the previous soil history, current agricultural management practices, and the plant-microbial community feedback mechanisms (Bouffaud *et al*., 2014). These processes interact to yield a number of edaphic conditions previously identified to promote oomycete growth.

For example, soils with excessive or insufficient nutrients for their local microbial communities are prone to oomycete outbreaks, as soil nutrient imbalance provides niche space for them (Löbmann *et al*., 2016; Rojas *et al*., 2017). Indicators of soil nutrient balance may include conductivity, cation exchange capacity (EC), total nitrogen, and total carbon (Löbmann *et al*., 2016; Rojas *et al*., 2017; Karppinen *et al*., 2020). Although none of the measured edaphic factors were particularly related to the communities observed in Trial 1, oomycete rhizosphere and roots communities with lentil soil histories were strongly associated with EC in Trial 2 (Fig. 3). Soil moisture is another key factor in promoting oomycete growth and is compounded by seeding into cool (< 16°C) soils (Hwang *et al*., 2015; Rojas *et al*., 2017; Karppinen *et al*., 2020). These conditions favour the release and chemotaxis of oomycete zoospores (Martin & Loper, 1999; Fawke *et al*., 2015). Therefore, we might have expected to observe a more dramatic change in the oomycete rhizosphere community between the wetter season of Trial 1 and the dry season of Trial 2.

The importance of soil chemistry may also be reflected in the results of our co-inertia analysis. This analysis illustrated that although the oomycete and bacterial data tended to have a significant relationship, neither community was particularly impactful in the roots nor the rhizosphere, and nor did any of the three soil histories influence their relationship (S11). Given that both of these microbial communities were derived from the same soil samples, they were more likely to experience the same edaphic factors. Therefore, the lack of obvious influences in the co-inertia analysis could be due to the oomycete and bacterial communities being similarly constrained by their common soil chemistry.

Microbes largely share basic biological reactions to abiotic factors, such as changes to pH, temperature, or water availability (Martiny *et al*., 2015). For example, bacteria, fungi, oomycetes, among others, require water for chemotaxis and locomotion, as well for maintaining turgor pressure (Martiny *et al*., 2015; Rojas & Huang, 2018). Cellular function requires the correct regulation of pressure, without which cells are unable to grow, divide, or move (Rojas & Huang, 2018). Although microbes have evolved a number of specialized tactics to regulate osmolarity, water stress, among other limitations imposed by soil chemistry, remain common constraints (Martiny *et al*., 2015). Therefore, the homogeneity we observed from the co-inertia analysis could reasonably be due to the oomycete and bacterial communities being similarly constrained by their common soil chemistry.

## 5. Conclusion

Oomycetes are major global phytopathogens, yet are understudied compared to other microbes. Here, we have shown for the first time the important role of soil history in structuring oomycete rhizosphere and root communities. We tested three different soil histories and found that none of the five planted *Brassicaceae* oilseed crops were able to restructure the oomycete communities the following year. We also took a novel approach in investigating how oomycete and bacterial communities may have structured one another. To our knowledge this is the first demonstration of the weak impact between the two microbial communities. Rather, the similarities between the two microbial communities may be due to being constrained by common edaphic factors. This study advances our understanding of how different agricultural practices can impact future microbial communities differently. Our results also highlight the need for continued monitoring of oomycete biodiversity and quantification.

## Acknowledgements

This work was supported by the Natural Sciences and Engineering Research Council of Canada, CRD Fund (Grant Number: CRDPJ 500507-16), Canola Council of Canada and Saskatchewan Pulse Growers, which are gratefully acknowledged. We thank Yantai Gan and Lee Poppy for setting up and managing field experiments at Swift Current. We also thank Stéphane Daigle for assistance in statistical analysis, Chih-Ying Lay, Jacynthe Masse, Chantal Hamel, and Simon Joly and for their helpful comments and discussions. Finally, AB would like to thank Morgan Botrel for her talents in collaborating on the graphical abstract, as well as Simon Morvan, Alexis Carteron, and the Quebec Centre for Biodiversity Science for their support and encouragement.

## Author Contributions

AJCB performed the qPCR experiment and assembled the mock community, prepared the samples for sequencing, analyzed the data, and wrote the manuscript with input from all co-authors. LB conducted field trials 1 and 2 and collected data; MSA & MH designed the experiment, supervised the work, contributed reagents, analytical tools, and revised the manuscript.

## Data Accessibility

Sequencing data and metadata are available at NCBI Bioproject under accession number: PRJNA849532.

## Supplementary Materials

### Supplementary Tables

**Table S1.** Nitrogen (N), phosphorous (P), potassium (K), and sulfur (S), available in the soil upon establishing the Test Phase, and the fertilizer applied, in the two-phase crop rotation experiment at Swift Current, SK. Adapted from Hossain *et al*., 2019.

| Swift Current Test Phase Plot Fertilization (N-P<sub>2</sub>O<sub>5</sub>-K<sub>2</sub>O-S kg ha<sup>-1</sup>) |  |  |  |
| --- | --- | --- | --- |
|  | Soil History | Nutrients Available <sup>a</sup> | Fertilizer Applied |
| Trial 1 | Chem-fallow | 37-34-646-22 | 48-7-0-10 |
|  | Lentil | 20-33-578-28 | 65-7-0-10 |
|  | Wheat | 18-31-511-19 | 68-7-0-10 |
| Trial 2 | Chem-fallow | 42-34-446-83 | 55-7-0-10 |
|  | Lentil | 30-43-488-83 | 43-7-0-10 |
|  | Wheat | 18-26-482-82 | 67-7-0-10 |

a, Measurements taken at *Brassicaceae* planting prior to fertilizing, with available N and S measured at 0–60 cm depth, P and K at 0–15 cm depth.

**Table S2.**
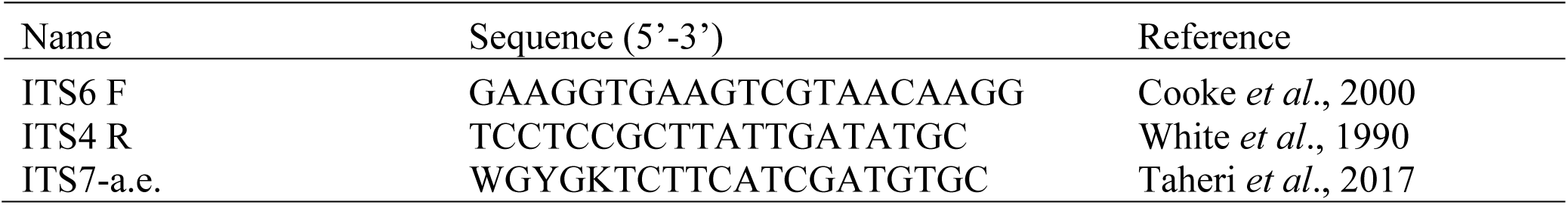
ITS primers used in this study.

**Table S3.** Oomycete strains that composed our mock community, which was included on each plate (Figure 2). The oomycete mock community we assembled contained DNA of 21 oomycete taxa in staggard copies / μL of ITS1 sequences, as per Bakker (2017). A copy of the mock community was included on each 96-well plate submitted for sequencing. Taxa are provided below to illustrate the level of comparison.

| Oomycetes |  | Copies of ITS |
| --- | --- | --- |
| Orders |  |  |
| Albuginales | *Albugo candida* 2v | ~ 100 |
| Saprolegniales | *Aphanomyces euteiches* 2 | 10 000 |
|  | *Aphanomyces euteiches* 206C | 1000 |
| Peronosporales | *Phytophthora* sp. 31594R | 1 000 000 |
|  | *Phytophthora* sp. 31461 | 500 000 |
|  | *Phytophthora* sp. 31710 | 250 000 |
|  | *Phytophthora* sp. 31598C | 100 000 |
| Pythiales | *Pythium* sp. 31309R | 20 000 000 |
|  | *Pythium* sp. 31411R | 1 000 000 |
|  | *Pythium* sp. 31392R | 1 000 000 |
|  | *Pythium* sp. 31799 | 750 000 |
|  | *Pythium* sp. 30717 | 500 000 |
|  | *Pythium* sp. 31019R | 500 000 |
|  | *Pythium* sp. 31800 | 250 000 |
|  | *Pythium* sp. 30918R | 100 000 |
|  | *Pythium* sp. 30623 | 100 000 |
|  | *Pythium* sp. 30886 | 75 000 |
|  | *Pythium ultimum* | 50 000 |
|  | *Pythium* sp. 30538 | 50 000 |
|  | *Pythium* sp. 30717 | 25 000 |
|  | *Pythium irregular* | 1000 |

### Supplementary Figures

**Figure S1.**
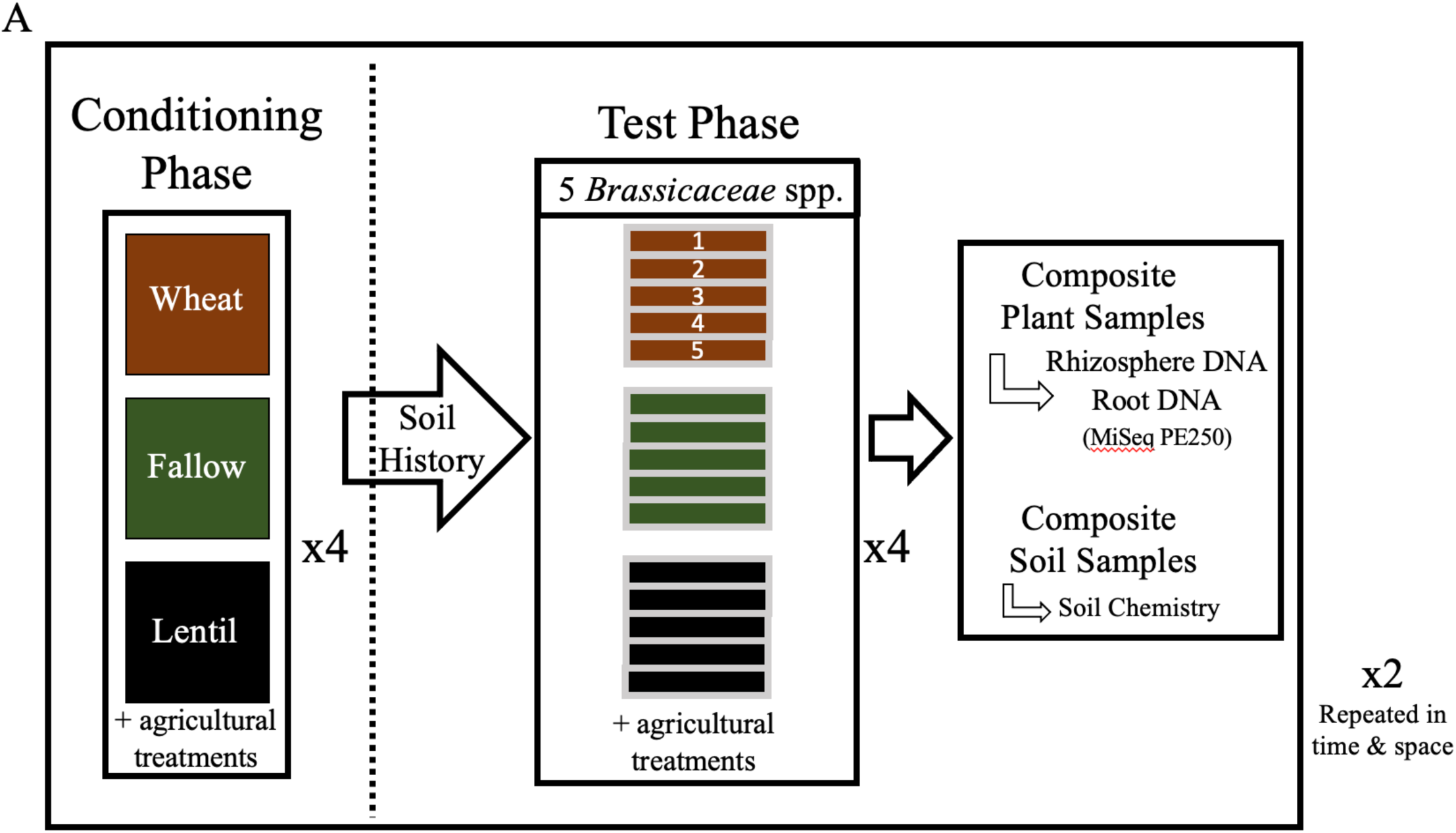
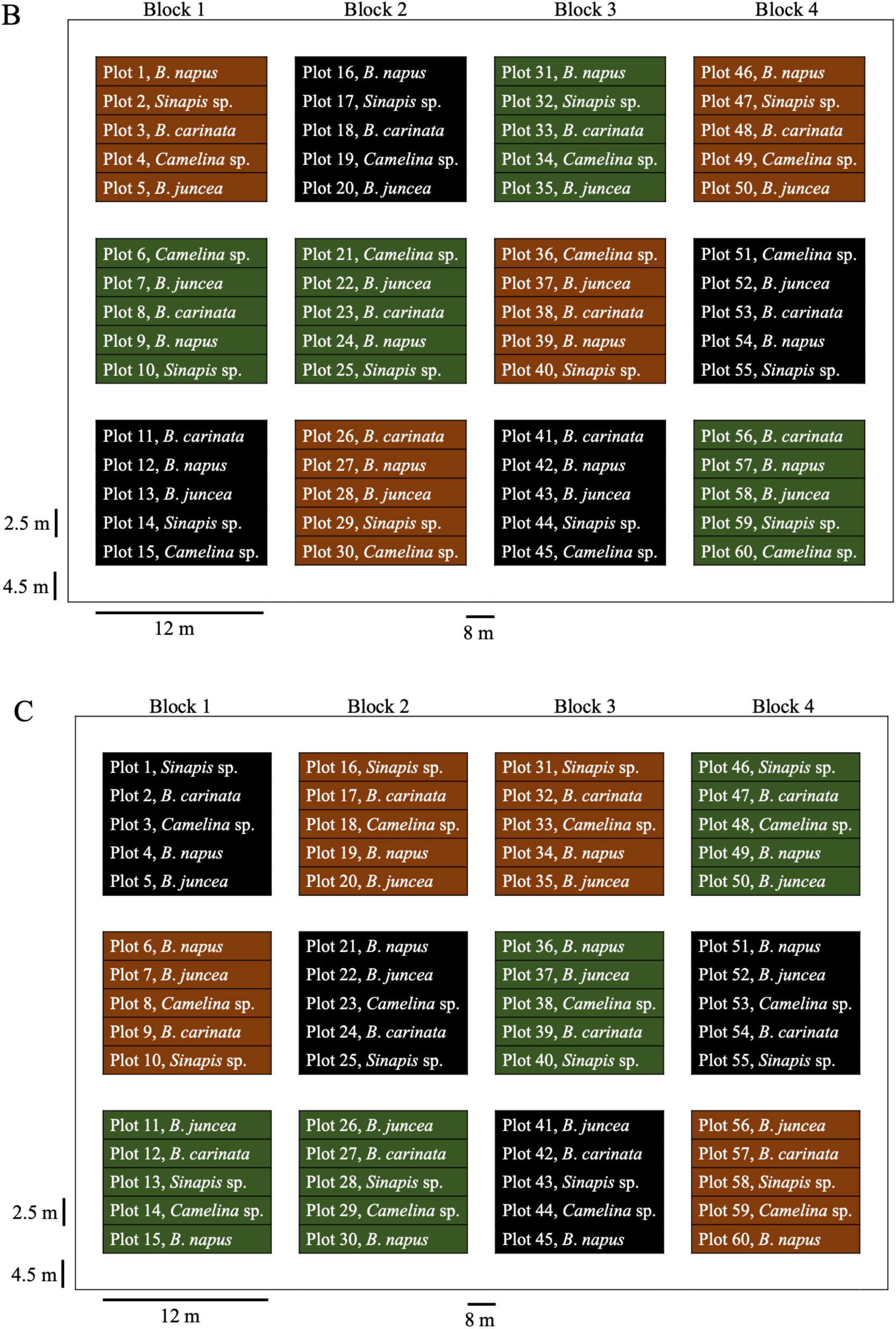
A two-year, or phase, cropping sequence was established in a field previously growing spring wheat (*Triticum aestivum* cultivar AC Lillian). The experimental design was a split-plot replicated in four complete blocks. In Phase 1, the ‘Conditioning Phase’, three soil history treatments were randomly assigned, consisting of spring wheat (*Triticum aestivum*, cv. AC Lillian), red lentil (*Lens culinaris* cv. CDC Maxim CL), or left fallow (brown, black, green, respectively). In Phase 2, the ‘Test Phase’, the conditioned plots were each subdivided and five *Brassicaceae* oilseed crop species were randomly assigned to one of these five subplots. Thus, each experiment had 60 subplots to sample. The experiment was performed twice, Trial 1, 2015-2016, and Trial 2, 2016-2017, on adjacent sites. (B) Trial 1 field plan for the *Brassicaceae* crops, which were Ethiopian mustard (*Brassica carinata* L., cv. ACC110), canola (*B*. *napus* L., cv. L252LL), oriental mustard (*B*. *juncea* L., cv. Cutlass), yellow mustard (*Sinapis alba* L., cv. Andante), and camelia (*Camelina sativa* L., cv. Midas). Boarder space between plots and blocks is in white. (C) Trial 2 field plan for the same *Brassicaceae* crops. For further details of this well-described experiment and its design, see Hossain *et al*. (2019), Liu *et al*. (2019), and Wang *et al*. (2020).

**Figure S2.**
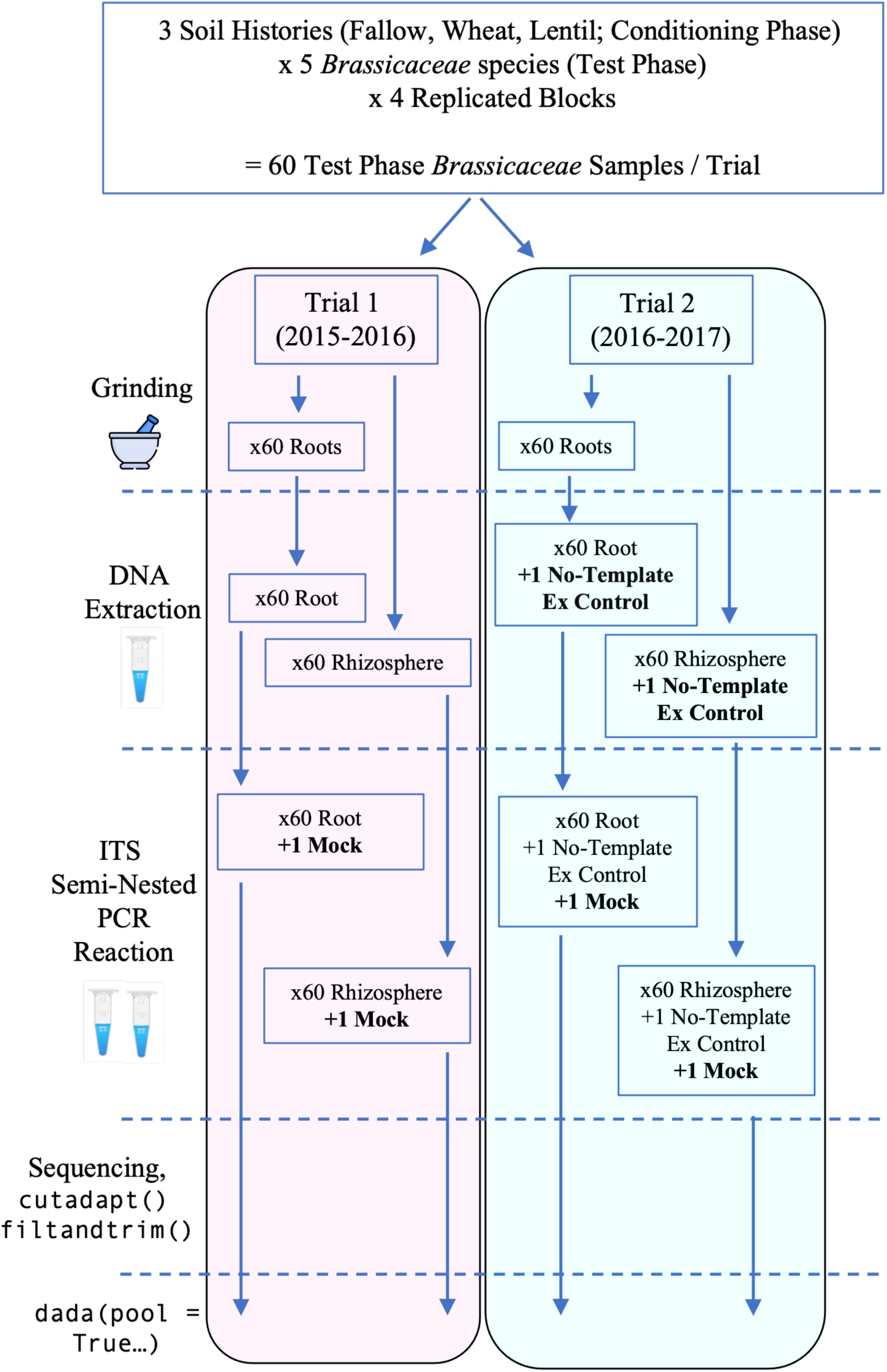
Organization of our lab workflow for the Test Phase *Brassicaceae* samples from harvest to generating amplicon sequence variants (ASVs). The Test Phase *Brassicaceae* samples were harvested in mid-late July. Four plants from two different locations within each of the 60 subplots were excavated and pooled together as a composite sample (Hossain *et al*., 2019; Liu *et al*., 2019, Wang *et al*., 2020). In the field, each plant had its rhizosphere soil divided from the root material, both portions were immediately flash-frozen in liquid nitrogen, and kept on ice. In the lab, roots were ground in liquid nitrogen, and DNA was extracted from all the Test Phase *Brassicaceae* root and rhizosphere portions. No-template extraction controls were included to assess what contaminates, or biases, the extraction kits might impart. We confirmed by gel electrophoresis that the no-template extraction controls contained DNA prior to sequencing. All DNA samples were submitted to Génome Québec for semi-nested ITS PCR amplification, library preparation, and paired-end 250 bp Illumina MiSeq sequencing. All reads were subsequently trimmed using cutadapt and processed through the DADA2 pipeline for ASV inference.

**Figure S3.**
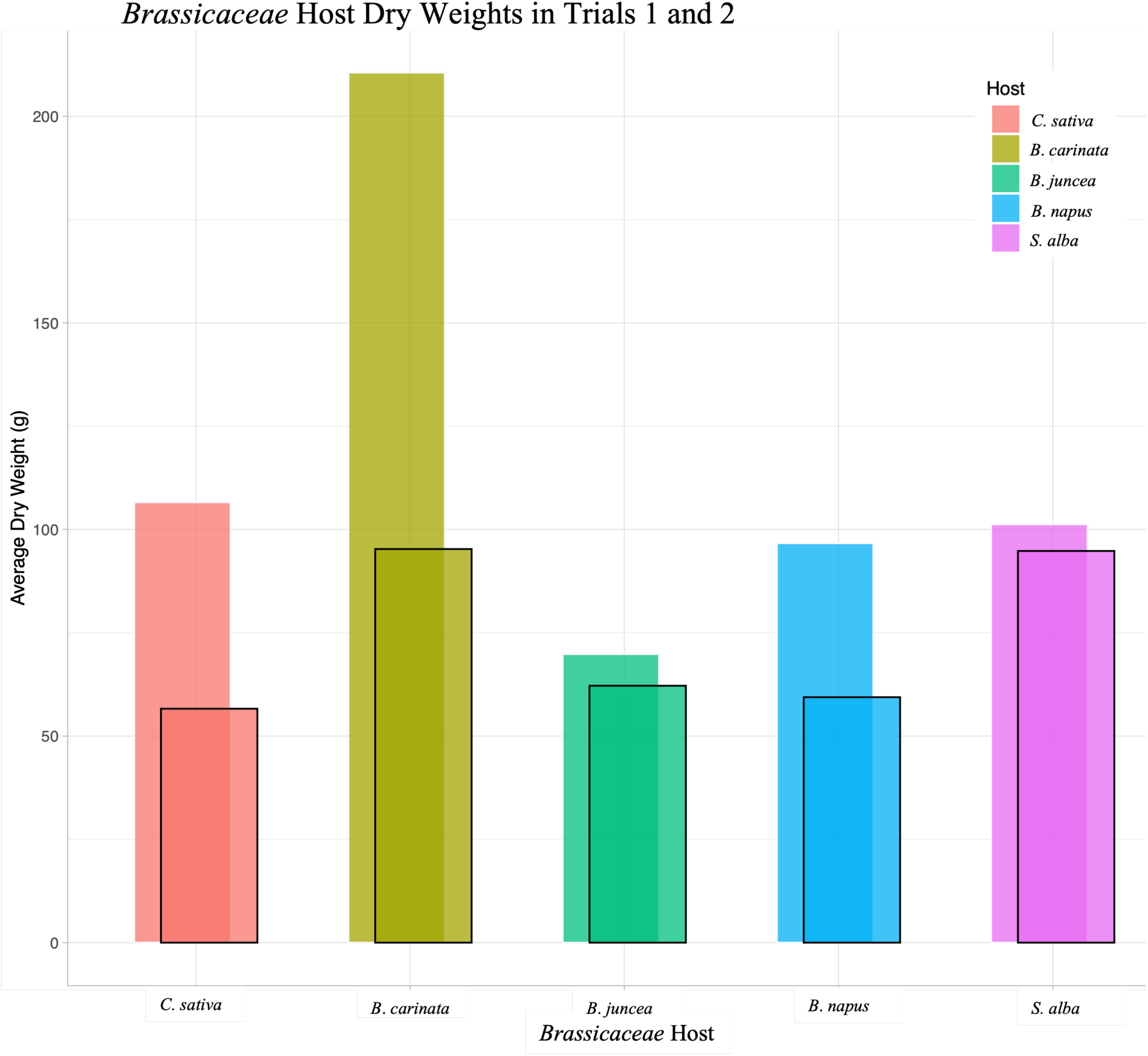
*Brassicaceae* host dry weights (g) decreased in Trial 2 (black outlines), compared to Trial 1 (no outlines). The Test Phase *Brassicaceae* samples were harvested in mid-late July, at Swift Current, Saskatchewan. The aerial portions were retained and dried to determine their weight.

**Figure S4.**
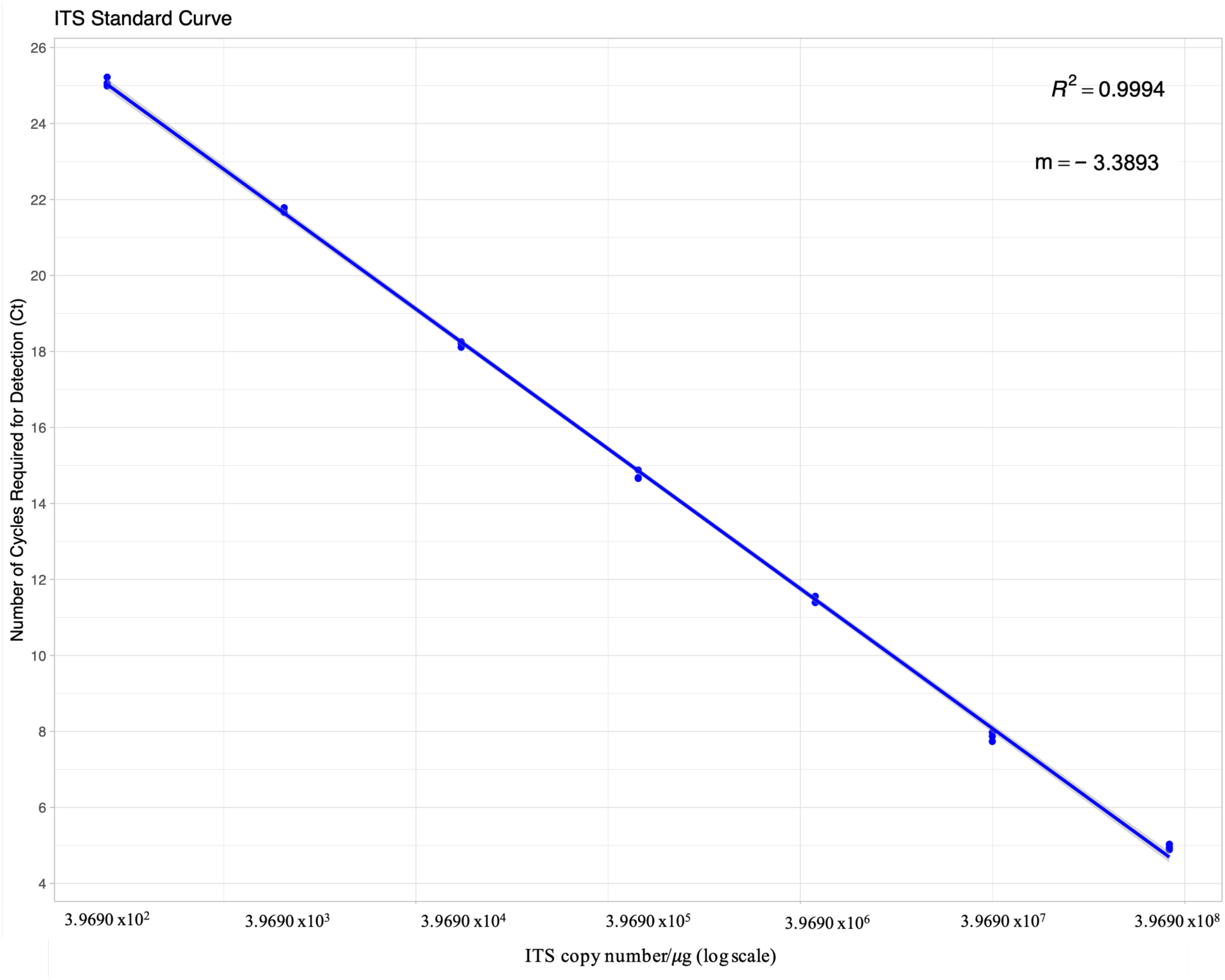
A standard curve of the ITS copy numbers (X-axis) versus the number of cycles required for detection (Ct, Y-axis), as determined from the serial dilution of a quantified ITS amplicon from concentrated *Pythium ultimum*.

**Figure S5.**
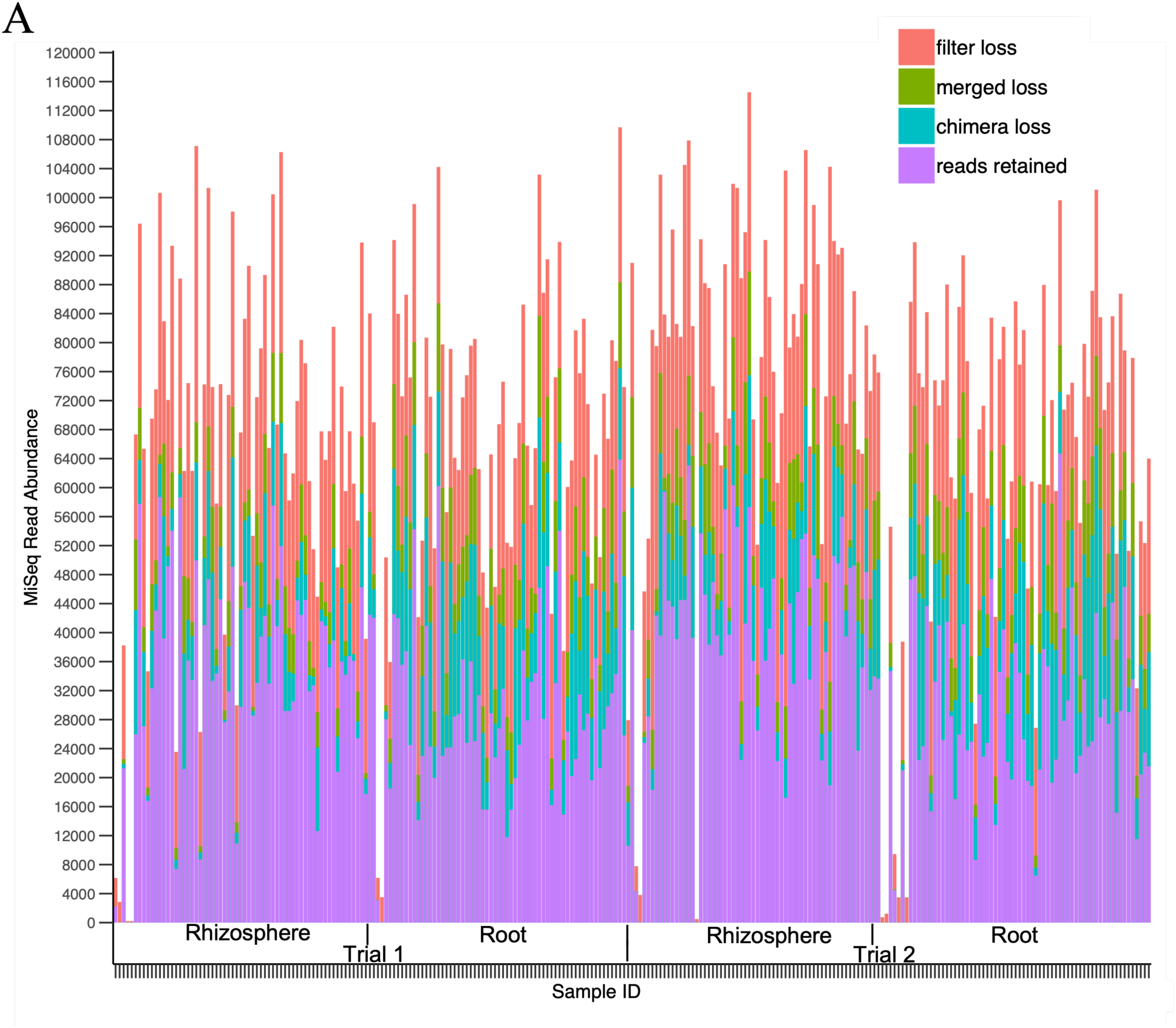
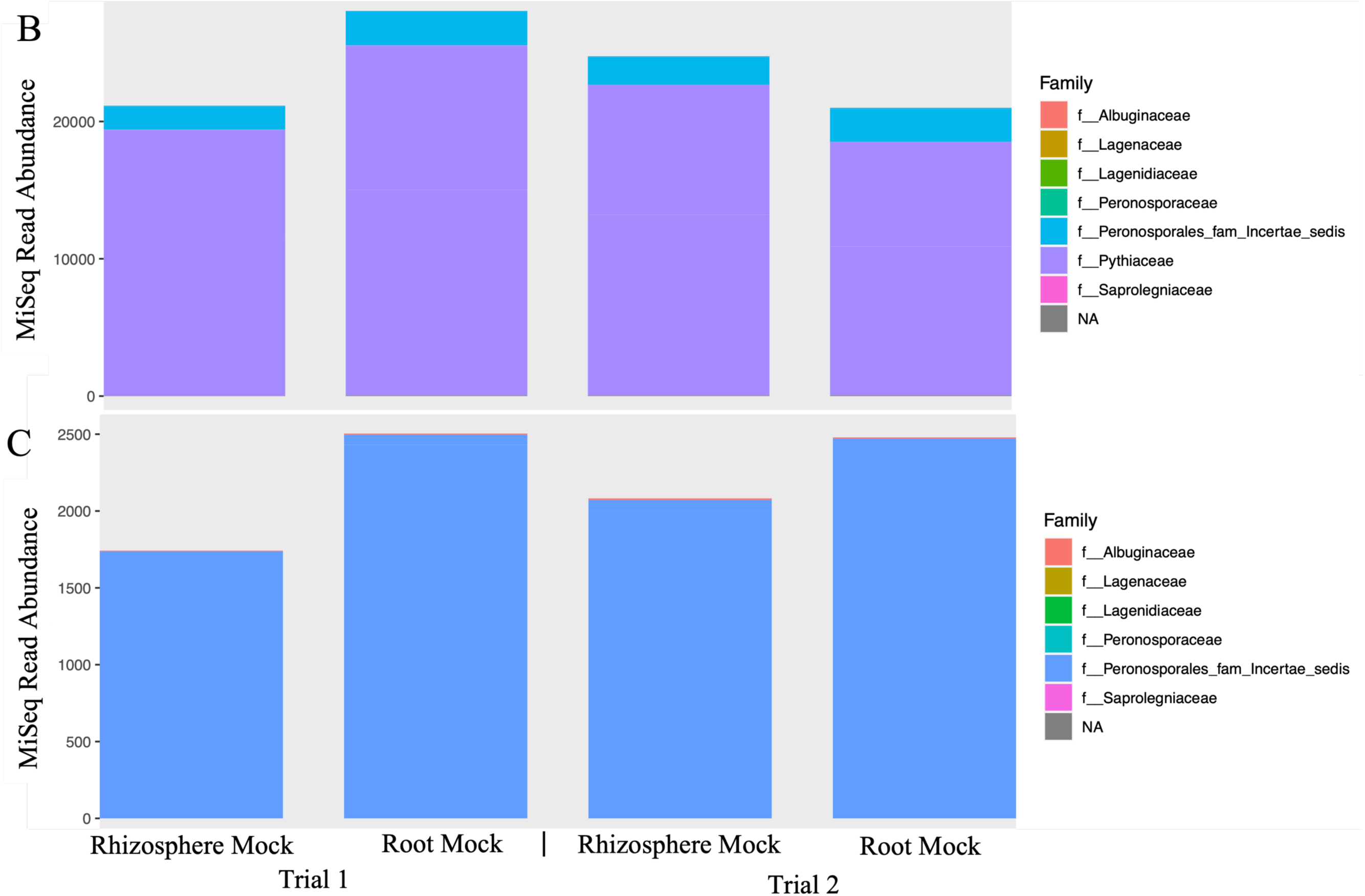
(A) The DADA2 workflow processed 17 656 076 raw reads produced from one lane of sequencing via Illumina’s MiSeq at Génome Québec in order to infer amplicon sequence variants (ASVs). Reads were produced in each of the no-template negative controls, although no DNA was detected in these samples post-extraction. The stringency of the `filterAndTrim` step eliminated nearly all the reads from these samples. 8 222 283 high-quality reads were retained among the Test Phase *Brassicaceae* samples. Among the Test Phase *Brassicaceae* samples, the rhizosphere samples retained noticeably more reads than their root partners. (B) Mock communities were assembled from known oomycete DNA and sequenced with experimental samples to confirm that various taxonomic groups of oomycetes were detectable with our pipeline. The mock communities were dominated by *Pythiaceae*, which is reflected in the composition of the mock sequences. (C) With the *Pythiaceae* sequences are removed, the number of *Peronosporales* and other groups become more evident.

**Figure S6.**
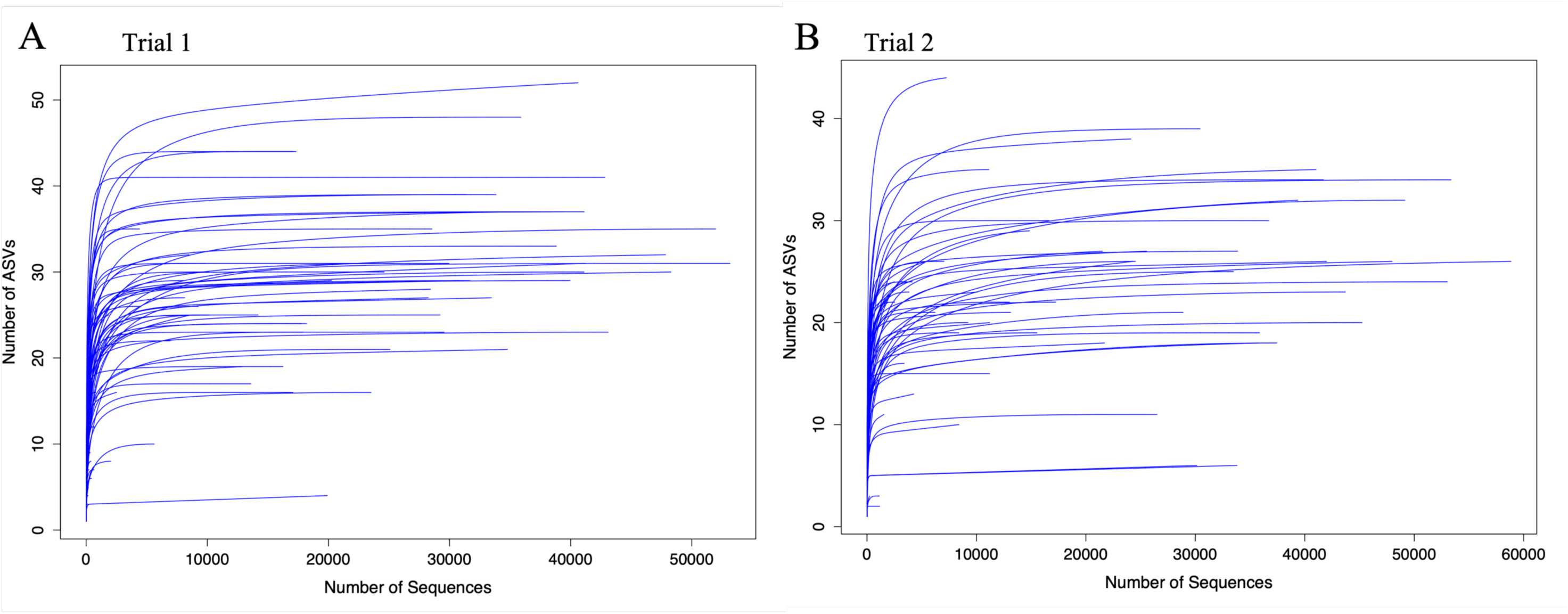
Rarefaction curves illustrated that the majority of the oomycete communities were identified in (A) Trial 1, and (B) Trial 2. The samples were harvested from two field trials during the Test Phase of a two-year crop rotation, in Swift Current, SK.

**Figure S7.**
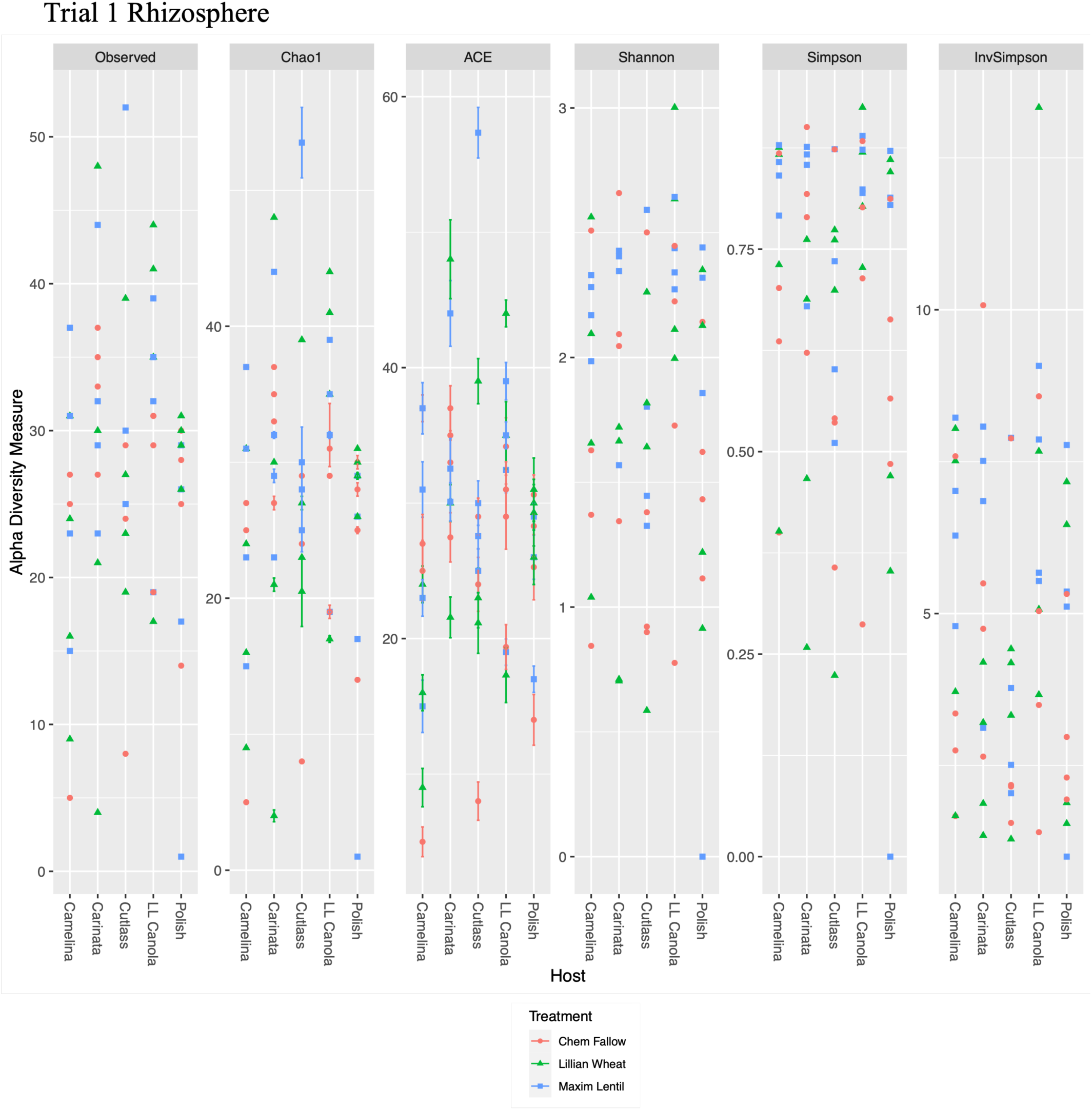
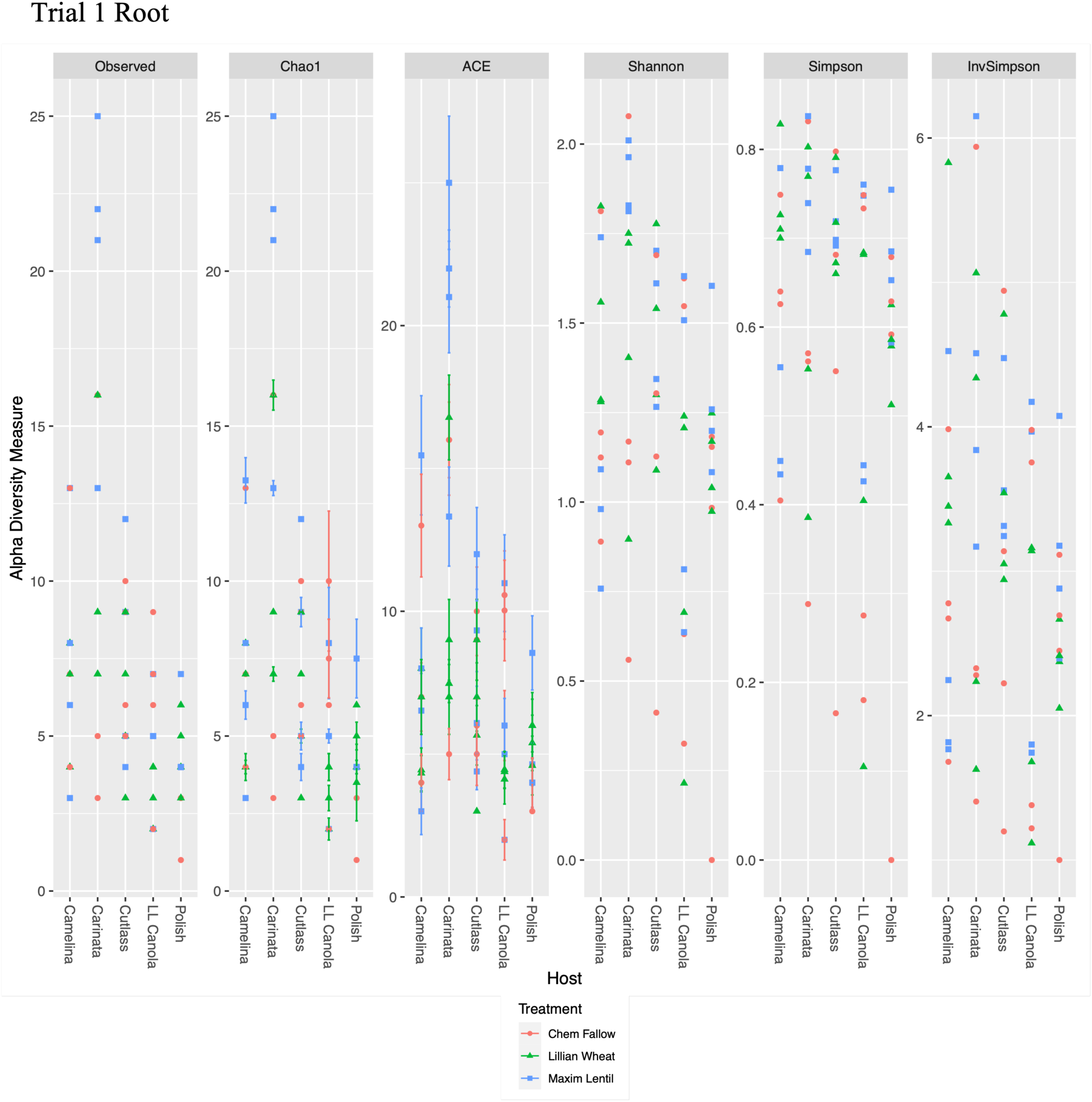
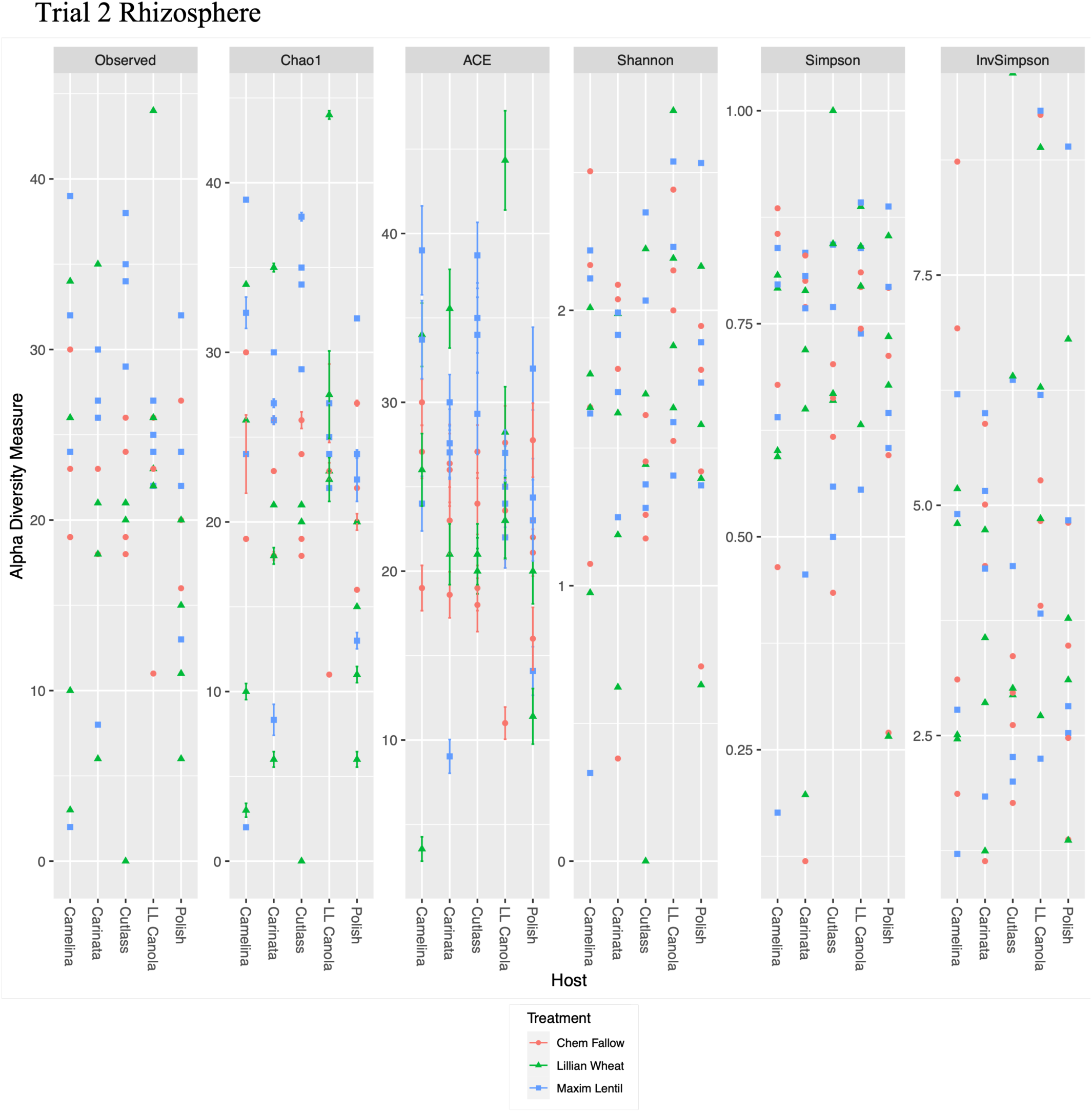
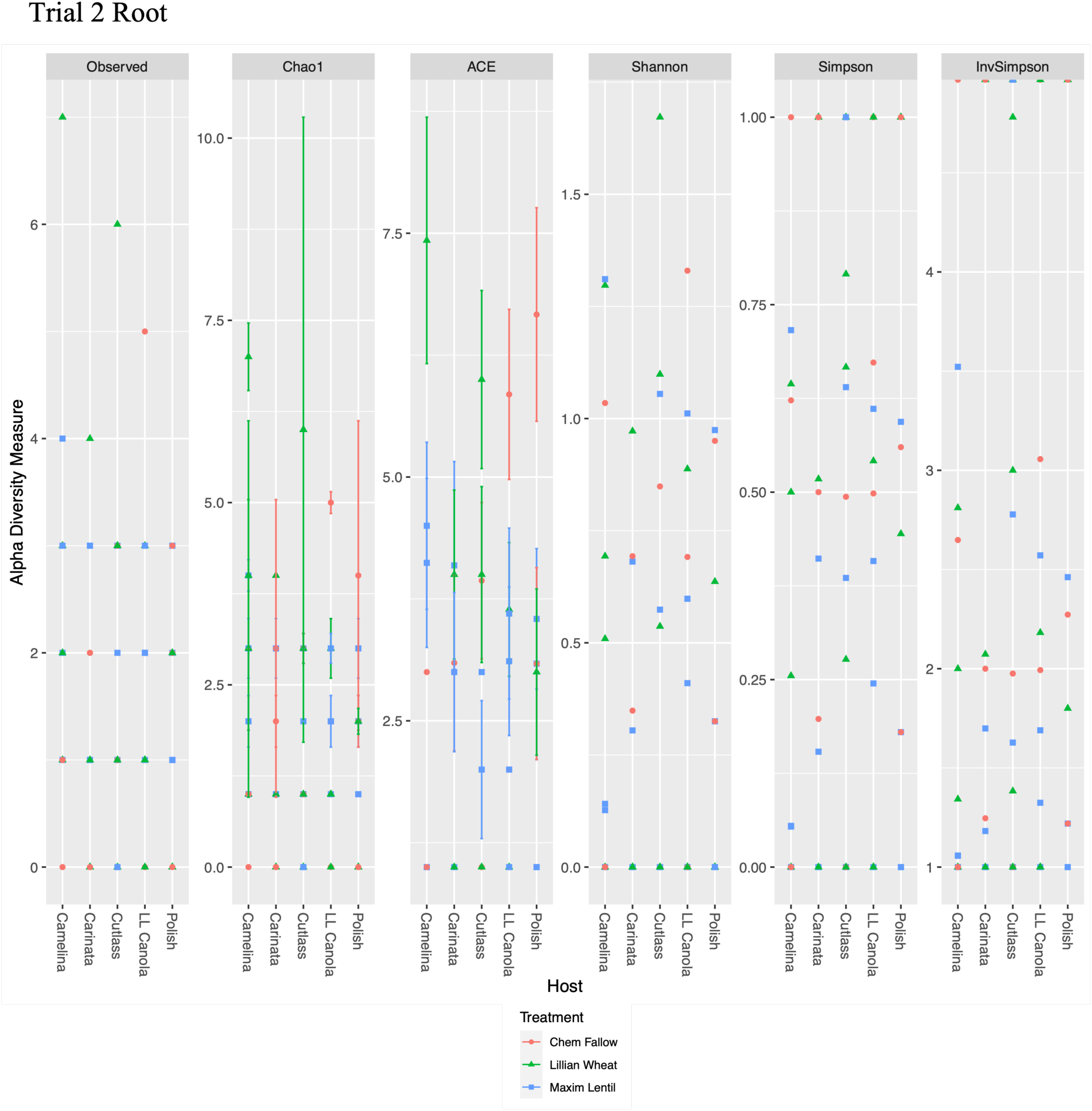
Taxa-based α-diversity indices (y-axis) for the rhizosphere (A & C) and root (B & D) communities from field trial 1, harvested 2016, and trial 2, harvest 2017. Each α-diversity index was grouped by *Brassicaceae* host, and reflect the phylogenetic diversity observed (Fig. 1), where communities are broadly similar across hosts, and soil histories.

**Figure S8.**
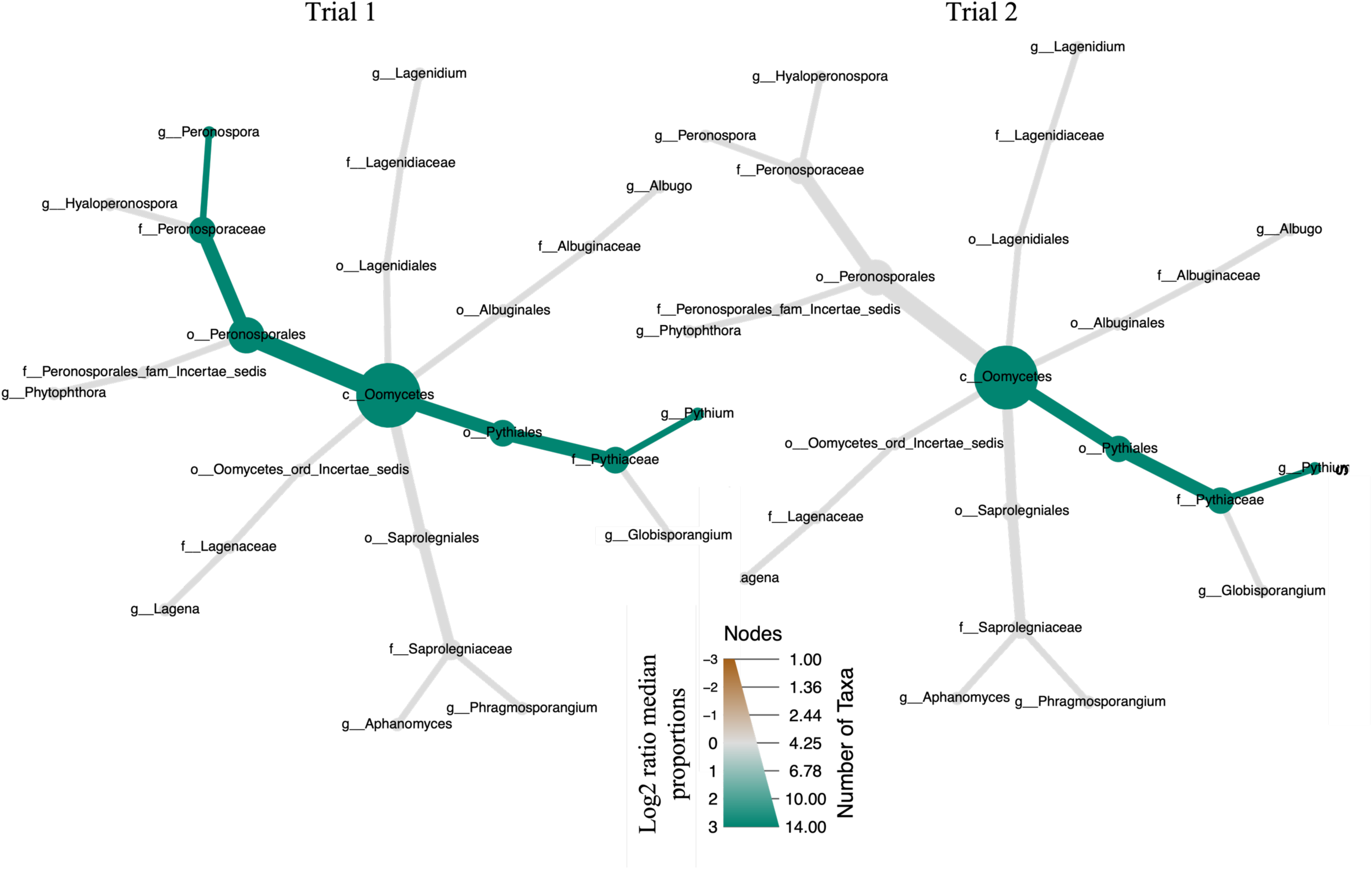
Differential taxa clusters of oomycete ASVs illustrated significantly more *Pythium* species in the rhizosphere communities compared to the roots in both field trials harvested from the Test Phase of a two-year rotation from Swift Current, Sask. *Peronospora* were also enriched in the rhizosphere communities in Trial 1, but not in Trial 2. Here, the size of the taxonomic groups (bubbles) represents the number of taxa, and the colour scale represents the proportion of each group, where the abundance of each taxonomic group in the cluster is compared between each compartment, using the using the non-parametric Kruskal test and the post-hoc pairwise Wilcox test, with the FDR correction. Taxa that are significantly (*p*. adj < 0.01) more abundant in the rhizosphere are highlighted in green.

**Figure S9.**
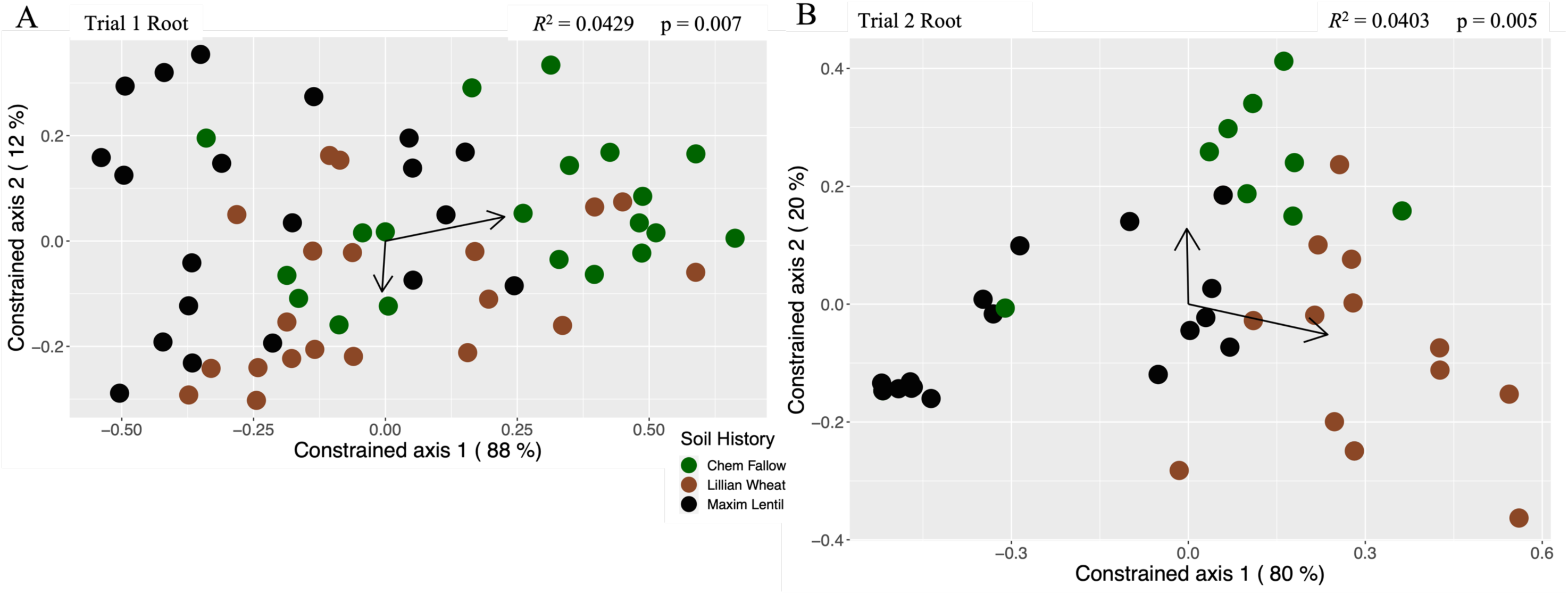
RDA illustrated that soil history structures the oomycete root communities in two field trials, Trial 1 (A, adj. *R^2^* = 0.0429, *p* = 0.007), and Trial 2 (B, adj. *R^2^* = 0.0403, *p* = 0.005), harvested from the Test Phase of a two-year rotation from Swift Current, SK.

**Figure S10.**
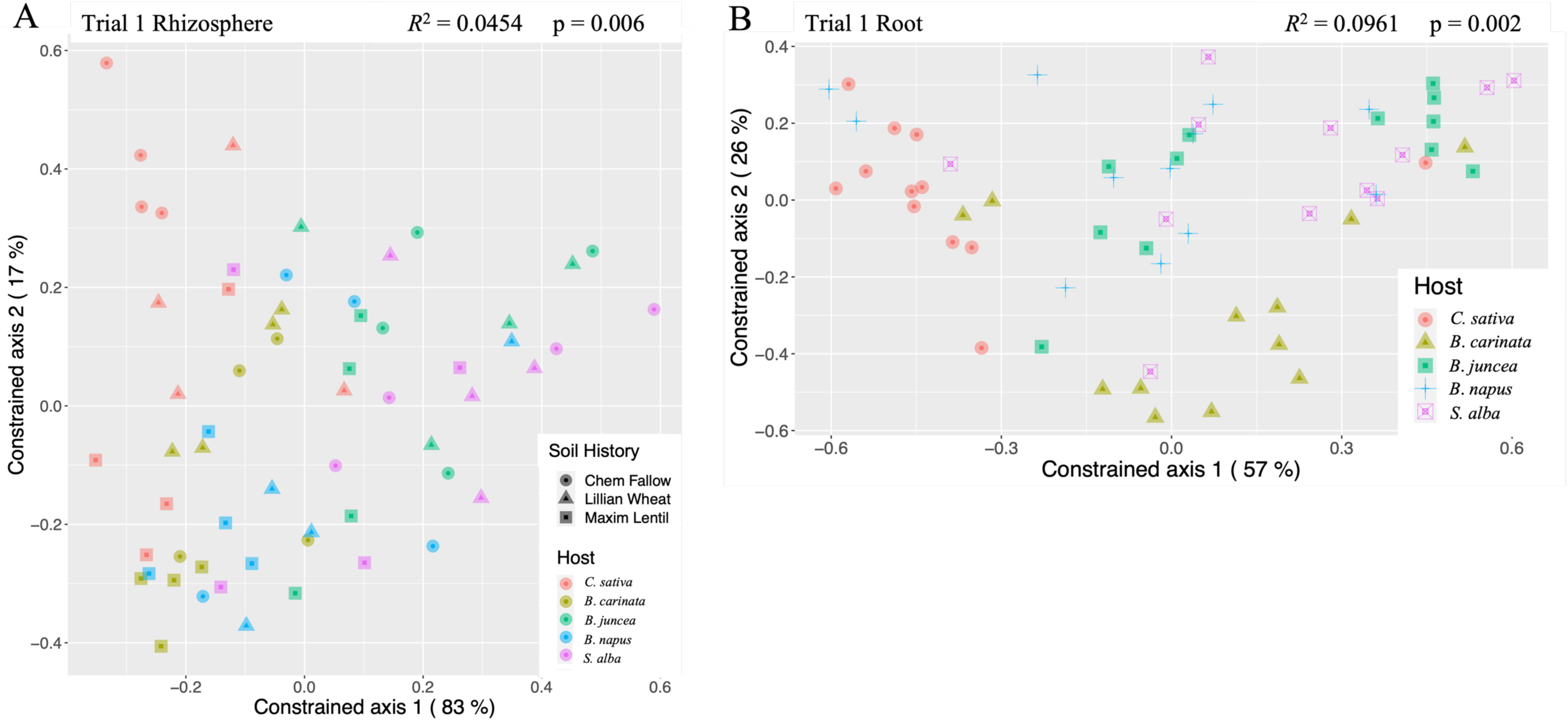
RDA illustrated the influence of *Brassicaceae* crop plants on the oomycete rhizosphere (A, adj. *R^2^*= 0.0454, *p* = 0.006) and root (B, adj. *R^2^* = 0.0961, *p* = 0.002) communities harvested from the Test Phase of field trial 1, as part of a two-year rotation from Swift Current, SK.

**Figure S11.**
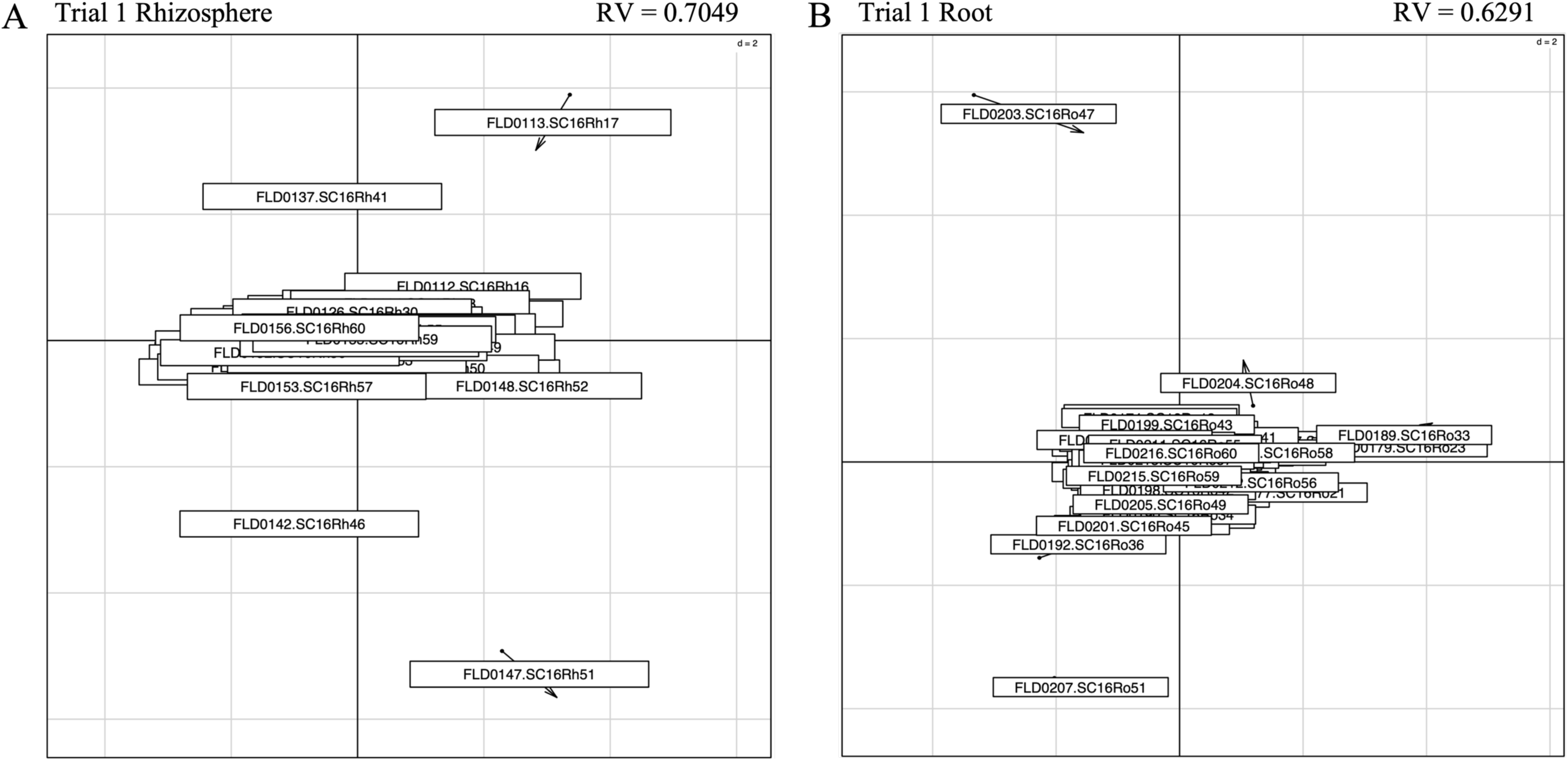
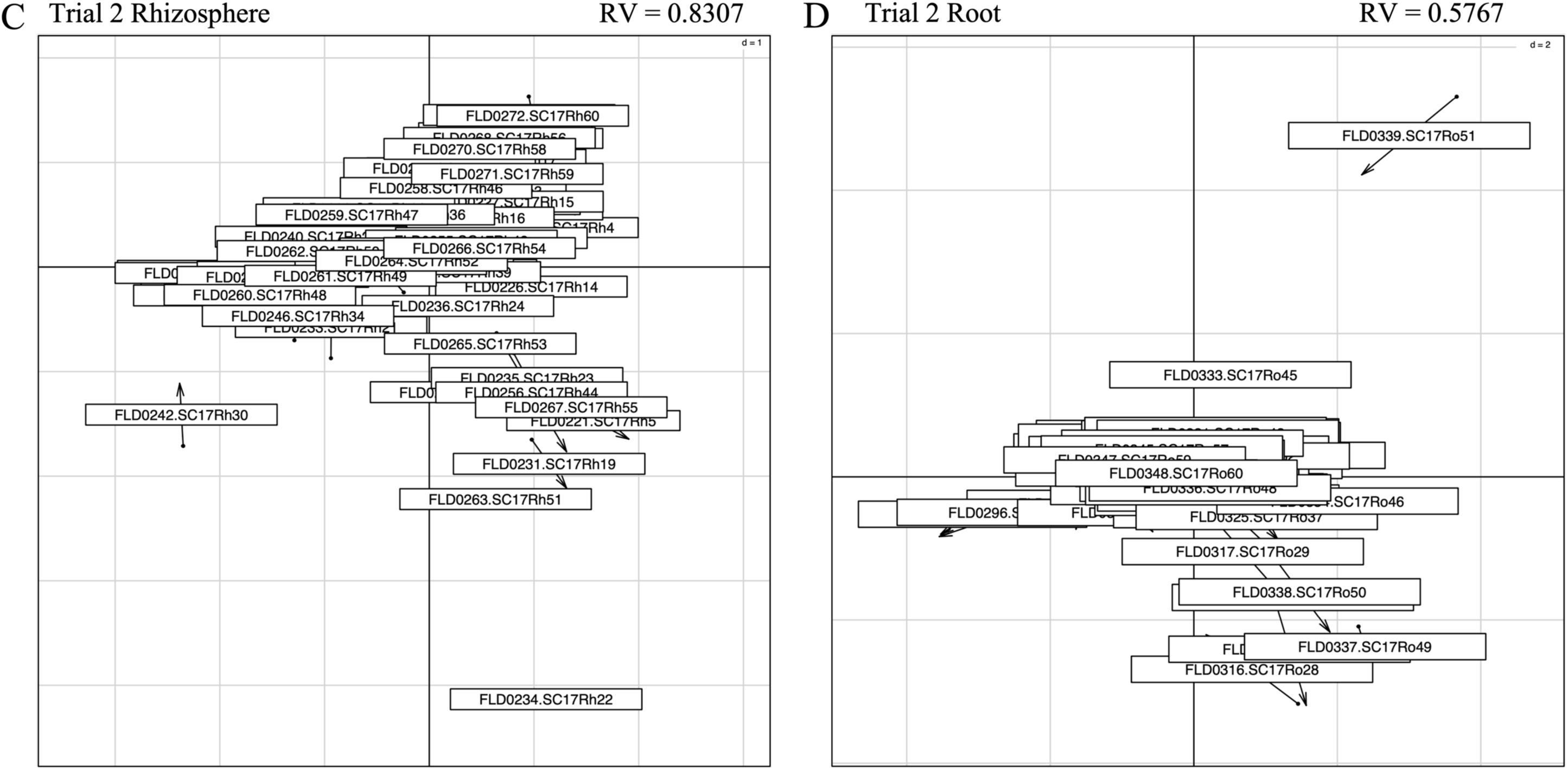
Co-inertia analysis illustrated that oomycete and bacterial communities were more significantly related in the rhizosphere (A, RV = 0.7049; C, RV = 0.8307) communities than their cognate root (B, RV = 0.6291; D, RV = 0.5767) samples harvested from the Test Phase of *Brassicaceae* field trial 1 (A & B), or trial 2 (C & D), as part of a two-year rotation from Swift Current, Sask. Moreover, the communities were largely similar, with very little divergence driven by any particular oomycete or bacterial ASV.

## References

Abarenkov K., Zirk A., Piirmann T., Pöhönen R., Ivanov F., Nilsson, R.H., & Kõljalg U. (2020) UNITE general FASTA release for eukaryotes. Version 04.02.2020. UNITE Community.

Bailey-Serres J., Parker J.E., Ainsworth E.A., Oldroyd G.E.D., & Schroeder J.I. (2019) Genetic strategies for improving crop yields. Nature 575:109–118.

Bakker M.G. (2018) A fungal mock community control for amplicon sequencing experiments. Molecular Ecology Resources 18:541–556.

Bakker P.A.H.M., Pieterse C.M.J., de Jonge R., & Berendsen R.L. (2021) The soil-borne legacy. Cell 172:1178–1180.

Bazghaleh N., Hamel C., Gan Y., Knight J.D., Vujanovic V., Cruz A.F., & Ishii T. (2016) Phytochemicals induced in chickpea roots selectively and non-selectively stimulate and suppress fungal endophytes and pathogens. Plant and Soil 409:479–493.

Bell T.H., Stefani F.O.P., Abram K., Champagne J., Yergeau É., Hijri M., & St-Arnaud M. (2016) A diverse soil microbiome degrades more crude oil than specialized bacterial assemblages obtained in culture. Applied Environmental Microbiology 82(18): 5530–5541.

Berendsen R.L., Vismans G., Yu K., Song Y., de Jonge R., Burgman W.P., Burmølle M., Herschend J., Bakker P.A.H.M., & Pieterse C.M.J. (2018) Disease-induced assemblage of a plant-beneficial bacterial consortium. ISME Journal 12:1496–1507.

Blakney A.J.C., Bainard L.D., St-Arnaud M., & Hijri M. (2022) *Brassicaceae* host plants mask the feedback from the previous year’s soil history on bacterial communities, except when they experience drought. Environmental Microbiology doi:10.1111/1462-2920.16046.

Borcard D., Legendre P., & Drapeau P. (1992) Partialling out the spatial component of ecological variation. Ecology 73: 1045–1055.

Bouffaud M.L., Poirier M. A., Muller D., & Moënne-Loccoz Y. (2014) Root microbiome relates to plant host evolution in maize and other *Poaceae*. Environmental Microbiology 16:2804–2814.

Callahan B.J., McMurdie P.J., & Holmes S.P. (2017) Exact sequence variants should replace operational taxonomic units in marker-gene data analysis. ISME Journal 11:2639–2643.

Callahan B.J., McMurdie P.J., Rosen M.J., Han A.W., Johnson A.J.A., & Holmes S.P. (2016a) DADA2: High-resolution sample inference from Illumina amplicon data. Nature Methods 13:581–583.

Callahan B.J., Sankaran K., Fukuyama J.A., McMurdie P.J., & Holmes S.P. (2016b) Bioconductor workflow for microbiome data analysis: from raw reads to community analyses. F1000Research 5:1492.

Canola Council of Canada, 2017. Canola Encyclopedia: Canola Growth Stages. Winnipeg, Manitoba, Canada.

Canola Council of Canada, 2017. Canola Encyclopedia: Diseases. Winnipeg, Manitoba, Canada.

Castrillo G., Teixeira P.J.P.L., Paredes S.H., Law T.F., de Lorenzo L., Feltcher M.E., Finkel O.M., Breakfield N.W., Mieczkowski P., Jones C.D., Paz-Ares J., & Dang J.L. (2017) Root microbiota drive direct integration of phosphate stress and immunity. Nature 543:513–518.

Cevik V., Boutrot F., Apel W., Robert-Seilaniantz A., Furzer O.J., Redkar A., Castel B., Kover P.X., Prince D.C., Holub E.B., & Jones J.D.G. (2019) Transgressive segregation reveals mechanisms of *Arabidopsis* immunity to *Brassica*-infecting races of white rust (*Albugo candida*). Proceedings of the National Academy of Science of the United States of America 116:2767–2773.

Chaparro J.M., Badri D.V., & Vivanco J.M. (2014) Rhizosphere microbiome assemblage is affected by plant development. ISME Journal 8:790–803.

Cooke D.E.L, Drenth A., Duncan J.M., Wagels G., & Brasier C.M. (2000) A molecular phylogeny of *Phytophthora* and related oomycetes. Fungal Genetics and Biology 30:17–32.

De Caceres M., & Legendre P. (2009) Associations between species and groups of sites: indices and statistical inference. Ecology 90(12): 3566–3574.

Delavaux C.S., Bever J.D., Karppinen E.M., & Bainard L.D. (2020) Keeping it cool: soil sample cold pack storage and DNA shipment up to 1 month does not impact metabarcoding results. Ecology Evolution 00:1–13.

Derevnina L., Petre B., Kellner R., Dagdas Y.F., Sarowar M.N., Giannakopoulou A., De la Concepcion J.C., Chaparro-Garcia A., Pennington H.G., van West P., & Kamoun S. (2016) Emerging oomycete threats to plants and animals. Philosophical Transactions of the Royal Society B 371:20150459.

Diéguez-Uribeondo J., García M.A., Cerenius L., Kozubíková E., Ballesteros I., Windels C., Weiland J., Kator H., Söderhäll K., & Martín M.P. (2009) Phylogenetic relationships among plant and animal parasites, and saprotrophs in *Aphanomyces* (Oomycetes). Fungal Genetics and Biology 45: 365–376.

Dolédec S., & Chessel D. (1994) Co-inertia analysis: an alternative method for studying species environment relationships. Freshwater Biology 31: 277–294.

Dray S., & Dufour A. (2007) The ade4 package: implementing the duality diagram for ecologists. Journal of Statistical Software 22 (4): 1–20.

Etesami H., & Alikhani H.A. (2016) Rhizosphere and endorhiza of oilseed rape (*Brassica napus* L.) plant harbor bacteria with multifaceted beneficial effects. Biological Control 94:1–24.

Fawke S., Doumane M., & Schornack S. (2015) Oomycete interactions with plants: infection strategies and resistance principles. Microbiology and Molecular Biology Reviews 79:263–281.

Fierer N., Jackson J.A., Vilgalys R., & Jackson R.B. (2005) Assessment of soil microbial community structure by use of taxon-specific quantitative PCR assays. Applied Environmental Microbiology 7(7): 4117–4120.

Fitzpatrick C.R., Copeland J., Wang P.W., Guttman D.S., Kotanen P.M., & Johnson M.T.J. (2018) Assembly and ecological function of the root microbiome across angiosperm plant species. Proceedings of the National Academy of Science of the United States of America: E1157–E1165.

Foster Z., Sharpton T., & Grünwald N. (2017) Metacoder: An R package for visualization and manipulation of community taxonomic diversity data. PLOS Computional Biology 13(2): 1–15.

Gavrin A., Rey T., Torode TA., Toulotte J., Chatterjee A., Kaplan J.L., Evangelisti E., Takagi H., Charoensawan V., Rengel D., Journet E.P., Debellé F., de Carvalho-Niebel F., Terauchi R., Braybrook S., & Schornack S. (2020) Developmental modulation of root cell wall architecture confers resistance to an oomycete pathogen. Current Biology 30: 4165–4176.

Godornes C., Leader B.T., Molini B.J., Centurion-Lara A., & Lukehart S.A. (2007) Quantitation of rabbit cytokine mRNA by real-time RT-PCR. Cytokine 38:1–7.

Gómez F.J.R., Navarro-Cerrillo R.M., Pérez-de-Luque A., Oβwald W., Vannini A., & Morales-Rodríguez C. (2021) Assessment of functional and structural changes of soil fungal and oomycete communities in holm oak declined dehesas through metabarcoding analysis. Scientific Reports 9:5315–5332.

Hamel C., Gan Y., Sokolski S., & Bainard L.D. (2018) High frequency cropping of pulses modifies soil nitrogen level and the rhizosphere bacterial microbiome in 4-year rotation systems of the semiarid prairie. Applied Soil Ecology 126:47–56.

Hannula S.E., Heinen R., Huberty M., Steinauer K., De Long J.R., Jongen R., & Bezemer T.M. (2021) Persistence of plant-mediated microbial soil legacy effects in soil and inside roots. Nature Communications 12:1–13.

Hossain Z., Johnson E.N., Wang L., Blackshaw R.E, & Gan Y. (2019) Comparative analysis of oil and protein content and seed yield of five *Brassicaceae* oilseeds on the Canadian prairie. Industrial Crop Production 136: 77–86.

Hou S., Thiergart T., Vannier N., Mesny F., Ziegler J., Pickel B., & Hacquard S. (2021) A microbiota–root–shoot circuit favours *Arabidopsis* growth over defence under suboptimal light. Nature Plants 7:1078–1092.

Hu L., Wu Z., Robert C.A.M., Ouyang X., Zust T., Mestrot A., Xu J., & Erb M. (2021) Soil chemistry determines whether defensive plant secondary metabolites promote or suppress herbivore growth. Proceedings of the National Academy of Science of the United States of America 118: e2109602118.

Hwang S.F., Ahmed H.U., Turnbull G.D., Gossen B.D., & Strelkov S.E. (2015) Effect of seeding date and depth, seed size and fungicide treatment on *Fusarium* and *Pythium* seedling blight of canola. Canadian Journal of Plant Science 95: 293–301.

Iffis B., St-Arnaud M., & Hijri M. (2016) Petroleum hydrocarbon contamination, plant identity and arbuscular mycorrhizal fungal (AMF) community determine assemblages of the AMF spore-associated microbes. Environmental Microbiology 18:2689–2704.

Kamoun S., Furzer O., Jones J.D.G., Judelson H.S., Ali G.S., Dalio R.J.D., Roy S.G., Schena L., Zambounis A., Panabières F., Cahill D., Ruocco M., Figueiredo A., Chen X.R., Hulvey J., Stam R., Lamour K., Gijzen M., Tyler B.M., Grünwald N.J. … (2015) The Top 10 oomycete pathogens in molecular plant pathology. Molecular Plant Pathology 16:413–434.

Kaisermann A., de Vries F.T., Griffiths R.I., & Bardgett R.D. (2017) Legacy effects of drought on plant–soil feedbacks and plant–plant interactions. New Phytologist 215: 1413–1424.

Karppinen E.M., Payment J., Chatterton S., Bainard J.D., Hubbard M., Gan Y., & Bainard L.D. (2020) Distribution and abundance of *Aphanomyces euteiches* in agricultural soils: effect of land use type, soil properties, and crop management practices. Applied Soil Ecology 150:103470.

Kawasaki A., Dennis P.G., Forstner C., Raghavendra A.K.H., Mathesius U., Richardson A.E., Delhaize E., Gilliham M., Watt M., & Ryan P.R. (2021) Manipulating exudate composition from root apices shapes the microbiome throughout the root system. Plant Physiology 187:2279–2295.

Kembel S.W., Cowan P.D., Helmus M.R., Cornwell W.K., Morlon H., Ackerly D.D., Blomberg S.P., & Webb C.O. (2010) Picante: R tools for integrating phylogenies and ecology. Bioinformatics 26: 1463–1464.

Korenblum E., Dong Y., Szymanski J., Panda S., Jozwiak A., Massalha H., Meir S., Rogachev I., & Aharoni A. (2020) Rhizosphere microbiome mediates systemic root metabolite exudation by root-to-root signaling. Proceedings of the National Academy of Science of the United States of America 117:3874–3883.

Krasnow C.S., & Hausbeck M.K. (2015) Pathogenicity of *Phytophthora capsici* to *Brassica* vegetable crops and biofumigation cover crops (*Brassica* spp.). Plant Disease 99:1721–1726.

Lau J.A., & Lennon J.T. (2012) Rapid responses of soil microorganisms improve plant fitness in novel environments. Proceedings of the National Academy of Science of the United States of America 109:14058–14062.

Lay C.Y., Bell T.H., Hamel C., Harker K.N., Mohr R., Greer C.W., Yergeau É., & St-Arnaud M. (2018) Canola root–associated microbiomes in the Canadian prairies. Frontiers in Microbiology 9:1–19.

Lebeis S.L., Paredes S.H., Lundberg D.S., Breakfield N., Gehring J., McDonald M., Malfatti S., del Rio T.G., Jones C.D., Tringe S.G., & Dangl J.L. (2015) Salicylic acid modulates colonization of the root microbiome by specific bacterial taxa. Science 349:860–864.

Legendre P., & De Cáceres M. (2013) Beta diversity as the variance of community data: dissimilarity coefficients and partitioning. Ecology Letters 16: 951–963.

Legendre P., & Legendre L. (2012) Numerical Ecology. Elsevier.

Liu B., Arlotti D., Huyghebaert B., & Tebbe C.C. (2022) Disentangling the impact of contrasting agricultural management practices on soil microbial communities – importance of rare bacterial community members. Soil Biology and Biochemistry 166:108573.

Liu K., Bandara M., Hamel C., Knight J.D., & Gan Y. (2020) Intensifying crop rotations with pulse crops enhances system productivity and soil organic carbon in semi-arid environments. Field Crop Research 248: 107657.

Liu K., Johnson E.N., Blackshaw R.E., Hossain Z., & Gan Y. (2019) Improving the productivity and stability of oilseed cropping systems through crop diversification. Field Crop Research 237: 65–73.

Löbmann M.T., Vetukuri R.R., de Zinger L., Alsanius B.W., Grenville-Briggs L.J., & Walter A.J. (2016) The occurrence of pathogen suppressive soils in Sweden in relation to soil biota, soil properties, and farming practices. Applied Soil Ecology 107:57–65.

Maciá-Vicente J.G., Nam B., & Thines M. (2020) Root filtering, rather than host identity or age, determines the composition of root-associated fungi and oomycetes in three naturally co-occurring *Brassicaceae*. Soil Biology and Biochemistry 146:107806.

Mamet S.D., Lamb E.G., Piper C.L., Winsley T., & Siciliano S.D. (2017) Archaea and bacteria mediate the effects of native species root loss on fungi during plant invasion. ISME Journal 11:1261–1275.

Marasco R., Rolli E., Ettoumi B., Vigani G., Mapelli F., Borin S., Abou-Hadid A.F., El-Behairy U.A., Sorlini C., Cherif A., Zocchi G., & Daffonchio D. (2012) A drought resistance-promoting microbiome is selected by root system under desert farming. PLoS One 7:e48479.

Martin F.N., & Loper J.E. (1999) Soilborne plant diseases caused by Pythium spp.: ecology, epidemiology, and prospects for biological control. Critical Reviews in Plant Science 18:111–181.

Martin M. (2011) Cutadapt removes adapter sequences from high-throughput sequencing reads. EMBnet.Journal 17(1): 10–12.

Martiny J.B.H., Jones S.E., Lennon J.T., & Martiny A.C. (2015) Microbiomes in light of traits: A phylogenetic perspective. Science 350: 1–8.

McMurdie P., & Holmes S. (2013) Phyloseq: an R package for reproducible interactive analysis and graphics of microbiome census data. PLoS One 8(4): e61217.

Mendes R., Kruijt M., de Bruijn I., Dekkers E., van der Voort M., Schneider J.H.M., Piceno Y.M., DeSantis T.Z., Andersen G.L., Bakker P.A.H.M., & Raaijmaker J.M. (2011) Deciphering the rhizosphere microbiome for disease-suppressive bacteria. Science 332:1097–1100.

Mohammed A.E., You M.P., Banga S.S., & Barbetti M.J. (2019) Resistances to downy mildew (*Hyaloperonospora brassicae*) in diverse *Brassicaceae* offer new disease management opportunities for oilseed and vegetable crucifer industries. European Journal of Plant Pathology 153:915–929.

O’Donovon J.T., Grant C.A., Blackshaw R.E., Harker K.N., Johnson E.N., Gan Y., Lafond G.P., May W.E., Turkington T.K., Lupwayi N.Z., Stevenson F.C., McLaren D.L., Khakbazan M., & Smith E.G. (2014) Rotational effects of legumes and non-legumes on hybrid canola and malting barley. Agronomy Journal 106: 1921–1932.

Oksanen J., Blanchet F.G., Friendly M., Kindt R., Legendre P., McGlinn D., Minchin P.R., O’Hara R.B., Simpson G.L., Solymos P., Stevens M.H.H., Szoecs E., & Wagner H. (2020). Vegan: Community Ecology Package. R package version 2.5-7

Preece C., Verbruggen E., Liu L., Weedon J.T., & Penuelas J. (2019) Effects of past and current drought on the composition and diversity of soil microbial communities. Soil Biology and Biochemistry 131:28–39.

Prince D.C., Rallapalli G., Xu D., Schoonbeek H.J., Çevik V., Asai S., Kemen E., Cruz-Mireles N., Kemen A., Belhaj K., Schornack S., Kamoun S., Holub E.B., Halkier B.A., & Jones J.D.G. (2017) *Albugo*-imposed changes to tryptophan-derived antimicrobial metabolite biosynthesis may contribute to suppression of non-host resistance to *Phytophthora infestans* in *Arabidopsis thaliana*. BMC Biology 15:20.

R Core Team (2020). R: A language and environment for statistical computing. R Foundation for Statistical Computing, Vienna, Austria.

Revillini D., Gehring C.A., & Johnson N.C. (2016) The role of locally adapted mycorrhizas and rhizobacteria in plant–soil feedback systems. Functional Ecology 30:1086–1098.

Richardson A.E., Barea J.M., McNeill A.M., & Prigent-Combaret C. (2009) Acquisition of phosphorus and nitrogen in the rhizosphere and plant growth promotion by microorganisms. Plant and Soil 321:305–339.

Rojas E.R., & Huang K.C. (2018). Regulation of microbial growth by turgor pressure. Current Opinion in Microbiology, 42:62–70.

Rojas J.A., Jacobs J.L., Napieralski S., Karaj B., Bradley C.A., Chase T., Esker P.D., Giesler L.J., Jardine D.J., Malvick D.K., Markell S.G., Nelson B.D., Robertson A.E., Rupe J.C., Smith D.L., Sweets L.E., Tenuta A.U., Wise K.A., & Chilvers M.I. (2017) Oomycete species associated with soybean seedlings in North America—Part II: diversity and ecology in relation to environmental and edaphic factors. Phytopathology 107:293–304.

Roy S., Coldren C., Karunamurthy A., Kip N.S., Klee E.W., Lincoln S.E., Leon A., Pullambhatla M., Temple-Smolkin R.L., Voelkerding K.V., Wang C., & Carter A.B. (2018) Standards and guidelines for validating next-generation sequencing bioinformatics pipelines. Journal of Molecular Diagnostics 20:4–27.

Sapkota R., & Nicolaisen M. (2015) An improved high throughput sequencing method for studying oomycete communities. Journal of Microbiological Methods 110:33–39.

Sapp M., Ploch S., Fiore-Donno A.M., Bonkowski M., & Rose L.E. (2018) Protists are an integral part of the *Arabidopsis thaliana* microbiome. Environmental Microbiology 20:30–43.

Semchenko M.., Leff J.W., Lozano Y.M., Saar S., Davison J.., Wilkinson A, Jackson B.G., Pritchard W.J., De Long J.R., Oakley S., Mason K.E., Ostle N.J., Baggs E.M., Johnson D., Fierer N., & Bardgett R.D. (2018) Fungal diversity regulates plant-soil feedbacks in temperate grassland. Scientific Advances 4:eaau4578.

Schimel J., Balser T.C., & Wallenstein M. (2007) Microbial stress-response physiology and its implications for ecosystem function. Ecology 88:1386–1394.

Schliep K.P. (2011) Phangorn: phylogenetic analysis in R. Bioinformatics 27:592–593.

Schwelm A., Badstöber J., Bulman S., Desoignies N., Etemadi M., Falloon R.E., Gachon C.M.M., Legreve A., Lukeš J., Merz U., Nenarokova A., Strittmatter M., Sullivan B.K., & Neuhauser S. (2017) Not in your usual Top 10: protists that infect plants and algae. Molecular Plant Pathology 19:1029–1044.

Sikes B.A., Cottenie K., & Klironomos J.N. (2009) Plant and fungal identity determines pathogen protection of plant roots by arbuscular mycorrhizas. Journal of Ecology 97:1274–1280.

Statistics Canada, Production of Principal Field Crops, November 2021

Taheri A.E., Chatterton S., Gossen B.D., & McLaren D.L. (2017) Degenerate ITS7 primer enhances oomycete community coverage and PCR sensitivity to *Aphanomyces* species, economically important plant pathogens. Canadian Journal of Microbiology 63:769–779.

Wang G., & Dunbrack R.L. (2004) Scoring profile-to-profile sequence alignments. Protein Science 13:1612–1626.

Wang L., Gan Y., Bainard L.D., Hamel C., St-Arnaud M., & Hijri M. (2020) Expression of N-cycling genes of root microbiomes provides insights for sustaining oilseed crop production. Environmental Microbiology 22: 4545–4556.

Weidner S., Koller R., Latz E., Kowalchuk G., Bonkowski M., Scheu S., & Jousset A. (2015) Bacterial diversity amplifies nutrient-based plant–soil feedbacks. Functional Ecology 29:1341–1349.

Wickham H. (2016) ggplot2: Elegant Graphics for Data Analysis. Springer-Verlag New York.

Wright E.S. (2016) Using DECIPHER v2.0 to analyze big biological sequence data in R. The R Journal 8:352–359.

Yang T., Lupwayi N., St-Arnaud M., Siddique K.H.M., & Bainard L.D. (2021) Anthropogenic drivers of soil microbial communities and impacts on soil biological functions in agroecosystems. Global Ecology Conservation 27:e01521.

Yu P., He X., Baer M., Beirinckx S., Tian T., Moya Y.A.T., Zhang X., Deichmann M., Frey F.P., Bresgen V., Li C., Razavi B.S., Schaaf G., von Wirén N., Su Z., Bucher M., Tsuda K., Goormachtig S., Chen X., & Hochholdinger F. (2021) Plant flavones enrich rhizosphere *Oxalobacteraceae* to improve maize performance under nitrogen deprivation. Nature Plants 7:481–499.

## References

Hossain Z., Johnson E.N., Wang L., Blackshaw R.E., & Gan Y. (2019) Comparative analysis of oil and protein content and seed yield of five *Brassicaceae* oilseeds on the Canadian prairie. Ind Crop Prod 136: 77–86.

Liu K., Johnson E.N., Blackshaw R.E., Hossain Z., & Gan Y. (2019) Improving the productivity and stability of oilseed cropping systems through crop diversification. Field Crop Res 237: 65–73.

Wang L., Gan Y., Bainard L.D., Hamel C., St-Arnaud M., & Hijri M. (2020) Expression of N-cycling genes of root microbiomes provides insights for sustaining oilseed crop production. Environ Microbiol 22: 4545–4556.

